# Decoding concept representations in aphasia after stroke

**DOI:** 10.64898/2026.04.07.717076

**Authors:** Jerry Tang, Carly Millanski, Allison Chen, Lisa D. Wauters, Jordyn Anders, Shilpa Shamapant, Stephen M. Wilson, Alexander G. Huth, Maya L. Henry

## Abstract

Many stroke survivors with aphasia struggle to map their thoughts into words or motor plans. Neuroprostheses that decode concept representations could help these individuals communicate by predicting the words, phrases, or sentences that they are struggling to produce. Here we decoded concept representations measured using functional magnetic resonance imaging (fMRI) from participants with different aphasia profiles. The decoders generated continuous word sequences that could describe the concepts that the participants were hearing about, seeing, or imagining. To forecast how this approach would generalize across the heterogeneity of aphasia profiles, we characterized how stroke affects the anatomical organization and information capacity of conceptual processing. Mapping how concepts are organized across the brain, we found that conceptual tuning during non-linguistic processing was largely consistent between the participants with aphasia and neurologically healthy participants. Comparing information processing between the participants with aphasia and neurologically healthy participants, we found that both groups processed similar amounts of non-linguistic information. Our findings indicate that concept representations can be largely spared in individuals with aphasia and demonstrate how these representations can be decoded to support communication.

## Introduction

Aphasia is a communication disorder caused by damage to the language regions of the brain. The most common cause of aphasia is stroke, and it is estimated that nearly a third of stroke survivors experience aphasia^1^. Strokes can affect multiple brain regions that serve different linguistic functions, so individuals with post-stroke aphasia can exhibit a variety of language production and comprehension impairments^2–4^. However, these individuals often perform well on non-linguistic tasks^5,6^ and report knowing what they want to say but not knowing how to say it^7^. This suggests that their conceptual knowledge is often spared even when their access to lexical or phonological knowledge is disrupted^8–10^. Consequently, neuroprostheses could help these individuals communicate by decoding spared concept representations into intended language output.

Concepts appear to be encoded in a widely distributed network of brain regions known as the semantic system^11^. To measure concept representations in neurologically healthy participants, we previously recorded brain activity using functional magnetic resonance imaging (fMRI) while each participant listened to many hours of narrative stories^12^. We then used voxelwise modeling to estimate how the activity in each brain region is related to semantic features of the stimulus words^13^. This produces an encoding model that can take any word sequence and predict how the participant’s brain would respond. To decode new brain responses, we generated word sequences to make the predicted responses under the encoding model as similar as possible to the actual responses^14^. This produces a natural language approximation of the concepts that the participant was thinking about.

While this semantic decoding approach can successfully approximate the meaning of perceived and imagined stimuli, it has previously only been demonstrated in neurologically healthy participants. Individuals with aphasia often have comorbidities that make it difficult for them to tolerate the many hours of scanning required to train decoders^15^. Moreover, these individuals often have language comprehension impairments that could make it difficult to train decoders on brain responses to linguistic stimuli^3^. To simultaneously address these barriers, we developed an approach for transferring semantic decoders across participants by aligning their brain responses using a small set of common stimuli^16,17^. Because concept representations are shared across cognitive processes, the common stimuli can be either linguistic or non-linguistic, making this transfer learning approach robust to language comprehension impairments^18–21^.

Here we used these approaches to decode concept representations in participants with post-stroke aphasia. We trained encoding models on brain responses from neurologically healthy reference participants, transferred them to the participants with aphasia, and used the aligned models to decode new brain responses. We found that the decoder predictions could describe the concepts that the participants with aphasia were hearing about, seeing, or imagining. Because semantic decoding requires spared concept representations, we assessed the generality of this approach by characterizing how stroke affects conceptual processing. We found that the anatomical organization and information capacity of conceptual processing were largely spared even in participants with extensive lesions and severe speech and language and impairments. This demonstrates the potential for semantic decoding to support individuals with post-stroke aphasia regardless of their lesion profiles, speech and language impairments, or intended concepts.

### Participants with aphasia

We enrolled three participants with post-stroke aphasia (**Figure 1**). P1 was a right-handed man who suffered a left intraparenchymal hemorrhagic stroke resulting from an arteriovenous malformation 8 years prior to participation in the study. P2 was a right-handed man who suffered a left middle cerebral artery infarction resulting from an embolism 2 years prior to participation. P3 was a right-handed woman who suffered bilateral middle cerebral artery infarctions resulting from bilateral pulmonary emboli 4 years prior to participation. We manually delineated lesions on T1-weighted images using ITK-SNAP. We characterized speech and language processing using the Quick Aphasia Battery^22^, auditory verbal comprehension using the Peabody Picture Vocabulary Test^23^, and conceptual processing using the Pyramids and Palm Trees Test^24^ (**Table 1**).

**Figure 1.**
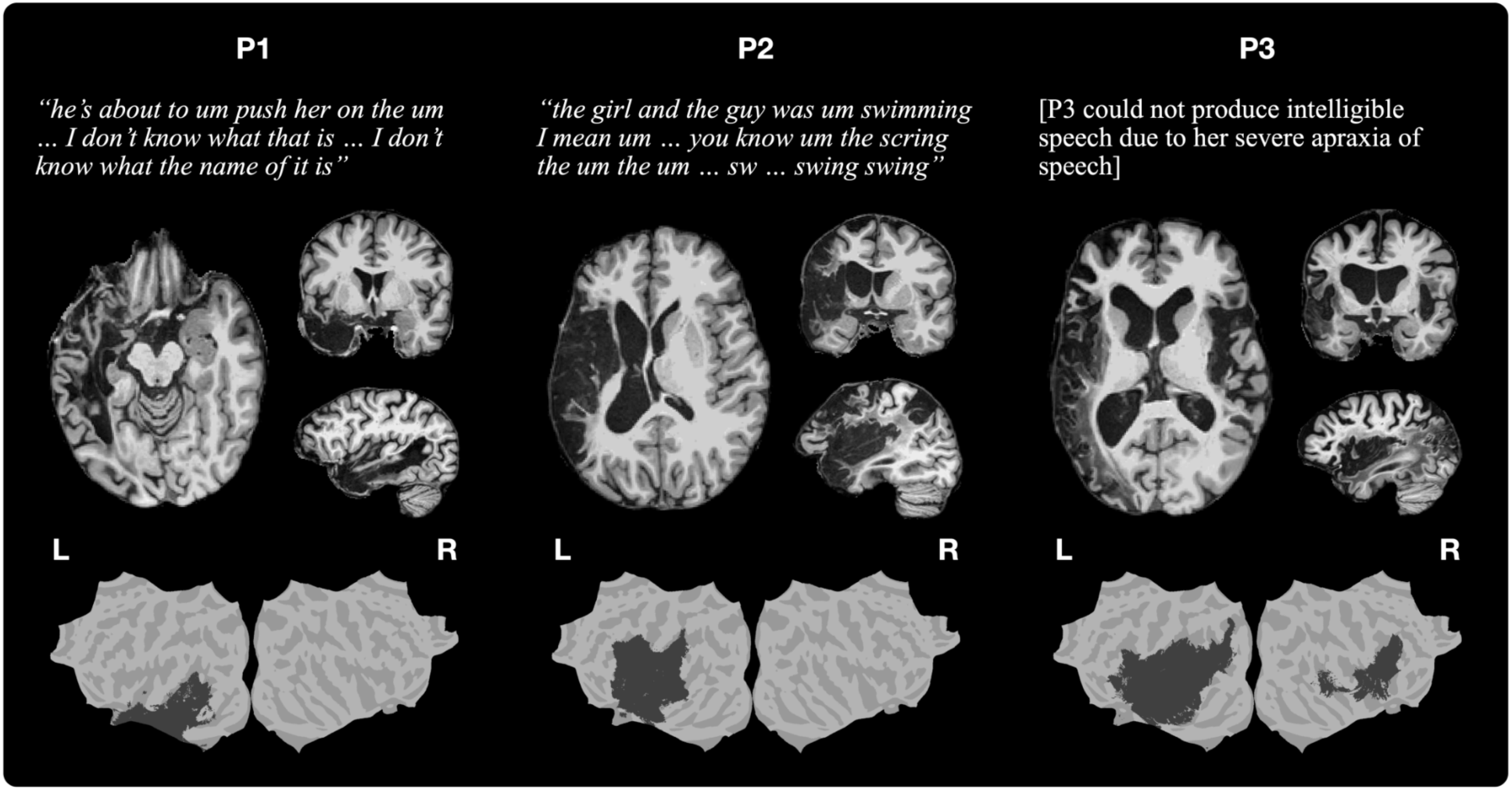
Participants with aphasia. Top: speech samples from P1 and P2 while they attempted to describe a picture of a boy pushing a girl on a swing. Middle: T1-weighted images from each participant. Bottom: cortical flatmaps with approximate lesion extent for each participant.

**Table 1.** Demographic information and behavioral assessment scores. Behavioral assessment was performed using the Quick Aphasia Battery^22^, Peabody Picture Vocabulary Test^23^, and Pyramids and Palm Trees–Pictures Test^24^. P1 had moderate aphasia predominantly affecting word finding and sentence comprehension. P2 had moderate aphasia predominantly affecting grammatical construction and word finding. P3 had severe expressive-receptive aphasia and severe apraxia of speech. Some assessment instructions were provided in writing for P3.

| Patient ID | P1 | P2 | P3 |
| --- | --- | --- | --- |
| **Demographic** |  |  |  |
| Age | 55 | 47 | 46 |
| Gender | Male | Male | Female |
| Education (years) | 17 | 13 | 17 |
| Handedness | Right | Right | Right |
| **Speech and language processing** |  |  |  |
| Quick Aphasia Battery: Auditory Word Comprehension (10) | 10 | 10 | 0.21 |
| Quick Aphasia Battery: Auditory Sentence Comprehension (10) | 1.67 | 6.67 | 3.75 |
| Quick Aphasia Battery: Written Word Comprehension (10) | 9.58 | 10 | 10 |
| Quick Aphasia Battery: Word Finding (10) | 4.75 | 4.75 | 1.75 |
| Quick Aphasia Battery: Grammatical Construction (10) | 8.25 | 1.63 | 0.50 |
| Quick Aphasia Battery: Writing (10) | 6.88 | 1.88 | 9.38 |
| Quick Aphasia Battery: Speech Motor Programming (10) | 10 | 10 | 0 |
| Quick Aphasia Battery: Repetition (10) | 9.17 | 5.83 | 1.67 |
| Quick Aphasia Battery: Reading Aloud (10) | 5.42 | 5.00 | 4.17 |
| Quick Aphasia Battery: Overall (10) | 6.83 | 6.30 | 1.65 |
| Peabody Picture Vocabulary Test (16) | 13 | 14 | 9 |
| **Conceptual semantic processing** |  |  |  |
| Pyramids and Palm Trees–Pictures (52) | 45 | 45 | 46 |

Structural imaging indicated that the participants had different lesion profiles. P1 had an extensive left hemisphere lesion impacting lateral and ventral temporal cortex. P2 had an extensive left hemisphere lesion impacting lateral frontal cortex, inferior parietal cortex, lateral temporal cortex, and insular cortex. P3 had an extensive left hemisphere lesion impacting lateral frontal cortex, inferior parietal cortex, lateral temporal cortex, and insular cortex, as well as a less extensive right hemisphere lesion impacting lateral frontal cortex, lateral temporal cortex, and insular cortex.

Speech and language testing indicated that the participants had impairments to different aspects of language production. P1 predominantly struggled with word finding; his spoken output was fluent with frequent pauses and circumlocutions. P2 struggled with grammatical construction and word finding; his spoken output was largely limited to short phrases and written output was largely limited to single words. P3 struggled with speech motor programming and grammatical construction; her spoken output was largely unintelligible due to severe apraxia of speech and written output was largely limited to short phrases. The participants also had different degrees of language comprehension impairment. P1 and P2 could comprehend conversational speech but struggled with more grammatically complex sentences, while P3 struggled to comprehend any spoken language but demonstrated much greater sparing of written language.

Despite their different speech and language impairments, participants showed relative sparing of nonverbal semantic processing on the Pyramids and Palm Trees–Pictures Test, a picture association task (45/52 for P1, 45/52 for P2, 46/52 for P3). The scores were close to the cutoff for neurologically healthy performance (47/52), indicating that non-linguistic conceptual knowledge was relatively spared. All participants demonstrated understanding of task instructions and complied with the behavioral assessments and neuroimaging experiments.

### Semantic decoding of perceived and imagined concepts

The goal of semantic decoding is to generate word sequences using only brain responses as input. To model the relationships between word sequences and brain responses, we trained encoding models on fMRI responses collected from four neurologically healthy reference participants while they listened to 10 h of narrative stories^12,13,25,26^ (**Figure 2a**). To transfer these encoding models to the participants with aphasia, we trained cross-participant converters to align fMRI responses between the reference participants and the participants with aphasia^16,17,27^. Converter training data collected from both groups comprised 3 h of narrative stories, as well as 1 h of silent movies to account for comprehension impairments. The participants with aphasia were given the option to listen to the stories with subtitles (**Methods**). To decode new brain responses from a participant with aphasia, we can generate word sequences using a language model, predict brain responses to the word sequences using the aligned encoding models, and identify the word sequences that are most consistent with the recorded brain responses^14^ (**Figure 2b**).

**Figure 2.**
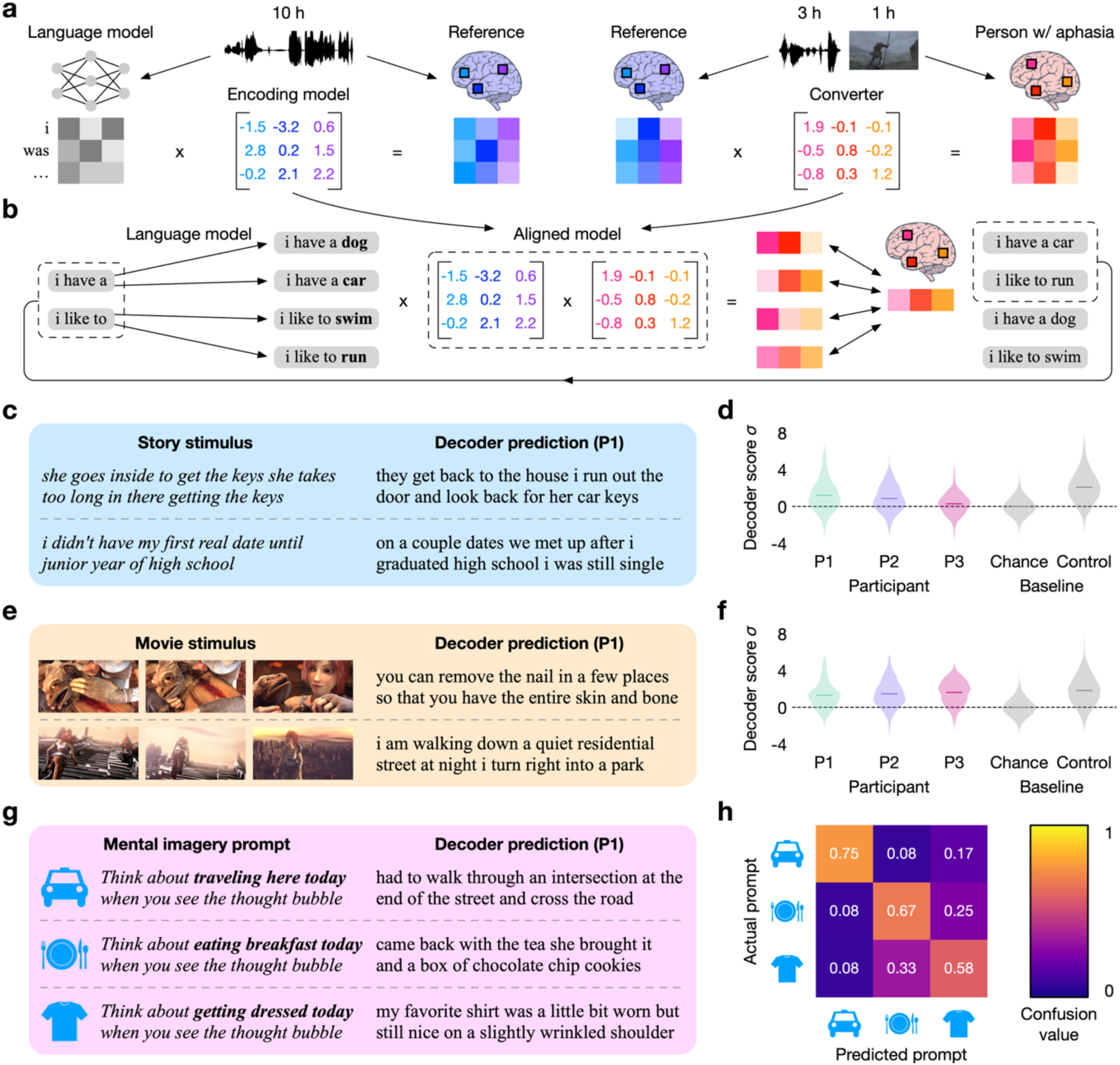
Semantic decoding of perceived and imagined concepts. **(a)** Encoding models were trained to predict brain responses collected from four neurologically healthy reference participants while they listened to 10 h of narrative stories. The encoding models were transferred to three participants with aphasia by aligning brain responses to 3 h of narrative stories and 1 h of silent movies. **(b)** To decode new brain responses from the participants with aphasia, a language model was used to generate word sequences and the aligned encoding models were used to identify the word sequences that were most consistent with the brain responses. **(c)** Decoder predictions from brain responses to test stories were compared to transcripts of the stories. **(d)** Story decoding performance for all participants with aphasia was significantly higher than expected by chance (q(FDR) < 0.05; one-sided non-parametric test) but lower than that of neurologically healthy controls. **(e)** Decoder predictions from brain responses to silent movies were compared to verbal descriptions of the movies. **(f)** Movie decoding performance for all participants with aphasia was significantly higher than expected by chance (q(FDR) < 0.05; one-sided non-parametric test) and approached that of neurologically healthy controls. **(g)** Decoder predictions from brain responses during mental imagery were compared to the prompted topics. **(h)** The decoder predictions could be used to identify the prompted topics with 67% accuracy (10/12 trials for P1; 8/12 for P2; 6/12 for P3; chance level 4/12).

To evaluate the semantic decoders, we first decoded single-trial brain responses collected while the participants listened to novel test stories. We found that the decoder predictions could successfully approximate the gist of the story events for P1 and P2 (**Figure 2c**; see **Supplementary Table 1** for all participants). We quantified decoding performance by comparing the predictions to the story words using BERTScore, which is a metric designed to measure the similarity of meaning between two word sequences^28^. Story decoding performance for all participants with aphasia was significantly higher than expected by chance (q(FDR) < 0.05; one-sided non-parametric test) but lower than that of neurologically healthy controls (**Figure 2d**). In particular, P3 had the lowest decoding performance consistent with her severe language comprehension impairment, which may have prevented her from accessing the conceptual meaning of the stories.

To determine whether the lower story decoding performance reflects conceptual processing impairments or language comprehension impairments, we next decoded single-trial brain responses collected while the participants watched novel test movies that were presented without sound. Our previous work showed that word sequences describing silent movie stimuli can be decoded from neurologically healthy participants using semantic decoders trained only on linguistic stimuli^14^. Here, we found that the decoder predictions could successfully approximate the gist of the movie events for all participants with aphasia (**Figure 2e**; see **Supplementary Table 2** for all participants). We quantified decoding performance by comparing the predictions to official verbal descriptions of the movies. Movie decoding performance for all participants with aphasia was significantly higher than expected by chance (q(FDR) < 0.05; one-sided non-parametric test) and approached that of neurologically healthy controls, indicating that non-linguistic conceptual processing was relatively spared (**Figure 2f**).

Language neuroprostheses should be able to decode internally generated concepts, so we decoded single-trial brain responses collected while the participants silently imagined three topics in the MRI scanner. We prompted the participants to imagine traveling to the scanning session, eating breakfast, and getting dressed. The participants then described what they had imagined for each topic outside of the scanner. We found that the decoder predictions often matched the prompted topics for all participants with aphasia (**Figure 2g**; see **Supplementary Table 3** for all participants). We quantified decoding performance by comparing the decoder predictions to the prompted topics and the participant descriptions. Mental imagery decoding performance for all participants with aphasia was significantly higher than expected by chance (q(FDR) < 0.05; one-sided non-parametric test) and the decoder predictions could be used to identify the prompted topics (24/36 trials; chance level 12/36) (**Figure 2h**). These results demonstrate the potential for language neuroprostheses to predict the concepts that individuals with aphasia are thinking about but struggling to produce.

### Anatomical organization of conceptual processing after stroke

The decoding results demonstrate that spared concept representations in the participants with aphasia can be decoded into continuous language. We can forecast how these results would generalize to other concepts from other individuals with aphasia by determining the extent to which concept representations are spared in stroke. We first mapped how stroke affects the anatomical organization of the semantic system. In neurologically healthy participants, concepts are encoded by distributed patterns of brain activity that are largely consistent across individuals^13^, stimulus modalities^20,21,29^, and languages^30^. One possibility is that spared brain regions in individuals with aphasia are selective for the same concepts before and after stroke, which would indicate that the semantic system can be robust to focal damage. Another possibility is that spared brain regions acquire selectivity for new concepts to compensate for damaged regions, which would indicate that the semantic system can reorganize in response to focal damage.

To evaluate these possibilities, we used encoding models to map the conceptual tuning of each brain region. We trained encoding models for the neurologically healthy reference participants using a lexical semantic embedding space derived from word co-occurrence statistics^13^. We transferred the encoding models to the participants with aphasia by aligning brain responses using either narrative stories or silent movies^17^. For each participant with aphasia, this produced a story-aligned encoding model that maps linguistic conceptual tuning and a movie-aligned encoding model that maps non-linguistic conceptual tuning. We then compared conceptual tuning between the participants with aphasia and the reference participants at every location on a cortical surface atlas^31^. Because the encoding models were transferred across participants using only functional correspondences, any anatomical correspondences between the encoding models indicate shared conceptual tuning.

To visualize conceptual tuning, we assigned a color to every location on the cortical surface atlas by projecting the encoding model weights onto the three dimensions of the lexical semantic embedding space that explained the most variance in a separate group of neurologically healthy participants^13^ (**Figure 3a**). The story-aligned encoding models showed that P1 shared linguistic tuning with the reference participants outside of damaged regions while P2 and P3 had larger differences in their linguistic tuning. However, the differences could result from impaired linguistic access to the concept representations rather than reorganization of the concept representations themselves. This was confirmed by the movie-aligned encoding models, which showed that all participants with aphasia shared non-linguistic tuning with the reference participants outside of damaged regions. For instance, concrete concepts (e.g. *visual*, *tactile*, *body part*) were represented near visual cortex and somatosensory cortex, while abstract concepts (e.g. *social*, *mental*, *time*) were represented in medial parietal cortex, inferior parietal cortex, and lateral temporal cortex^13^.

**Figure 3.**
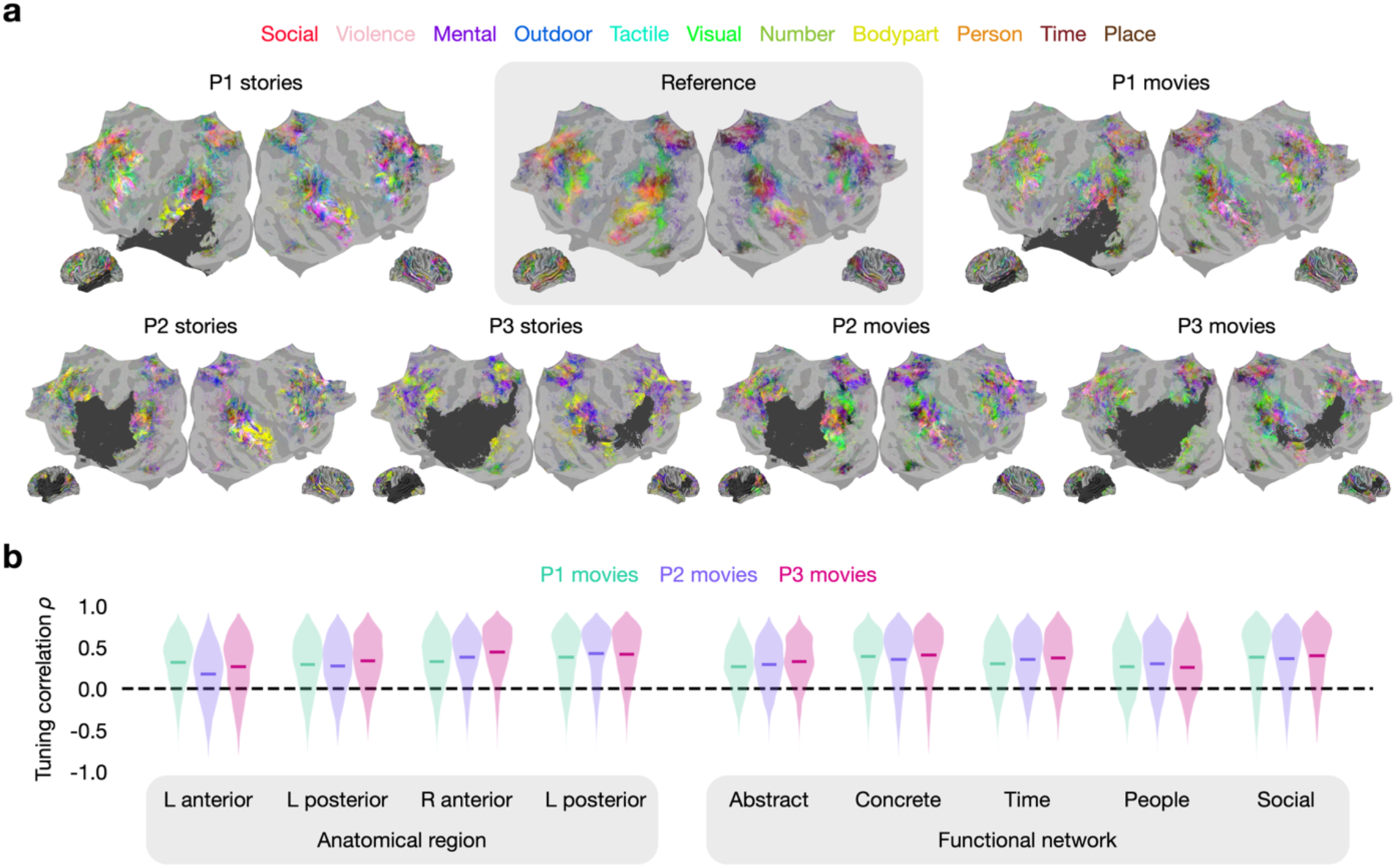
Anatomical organization of conceptual processing after stroke. **(a)** Encoding models were transferred from neurologically healthy reference participants to participants with aphasia by aligning brain responses using narrative stories or silent movies. Concept-selective cortical locations were colored based on their conceptual tuning. During linguistic processing, P1 shared tuning with the reference participants outside of damaged regions. During non-linguistic processing, all participants with aphasia shared tuning with the reference participants outside of damaged regions. **(b)** Non-linguistic tuning was compared between the participants with aphasia and the reference participants at every location on a cortical surface atlas. Shared tuning was quantified using the rank correlation between simulated brain responses to 10,470 concepts. Tuning was significantly more similar than expected by chance (q(FDR) < 0.05; one-sided permutation test) in all tested anatomical regions and functional networks.

To quantify these results, we operationalized the conceptual tuning at every location on the cortical surface atlas by using the movie-aligned encoding models to simulate brain responses to 10,470 concepts^13^. We then computed the shared tuning between each participant with aphasia and the reference participants using the rank correlation between the simulated brain responses. High tuning correlation in a brain region indicates that the region is selective for similar concepts in the participant with aphasia and the reference participants. Low tuning correlation indicates that the region is selective for different concepts. We found that tuning correlation was significantly higher than expected by chance (q(FDR) < 0.05; one-sided permutation test) in all tested anatomical regions and functional networks (**Figure 3b**). These results demonstrate that the anatomical organization of the semantic system can be robust to stroke, which is consistent with previous findings that concepts are redundantly encoded in multiple brain regions^13,14^. Consequently, damage to a brain region may not prevent us from decoding the concepts encoded in the region as long as the other regions that encode the concepts are spared.

### Information capacity of conceptual processing after stroke

We next measured how stroke affects the information capacity of the semantic system. Even if each brain region processes the same concepts after stroke as it did before stroke, the semantic system as a whole might not process information to the same extent (e.g. only processing concepts from certain categories) or granularity (e.g. only processing more superordinate concepts). Because semantic decoders operate on spared concept representations, the information capacity of the semantic system determines the extent and granularity of the concepts that can be decoded.

While we cannot measure the premorbid capacity of the semantic system in a participant with aphasia, we can approximate it using brain responses from neurologically healthy reference participants. We can then quantify the post-stroke capacity in the participant with aphasia by assessing how well their brain responses can predict brain responses from the reference participants. Here we separately trained cross-participant converters on brain responses to narrative stories and silent movies, and evaluated converter performance using the linear correlation between the predicted and actual responses from the reference participants (**Figure 4a**). High converter performance in a reference participant brain region indicates that the information processed in the region is still processed somewhere in the participant with aphasia^32^. Low converter performance indicates that the information may no longer be processed anywhere in the participant with aphasia. We obtained a ceiling on converter performance by training cross-participant converters between pairs of reference participants.

**Figure 4.**
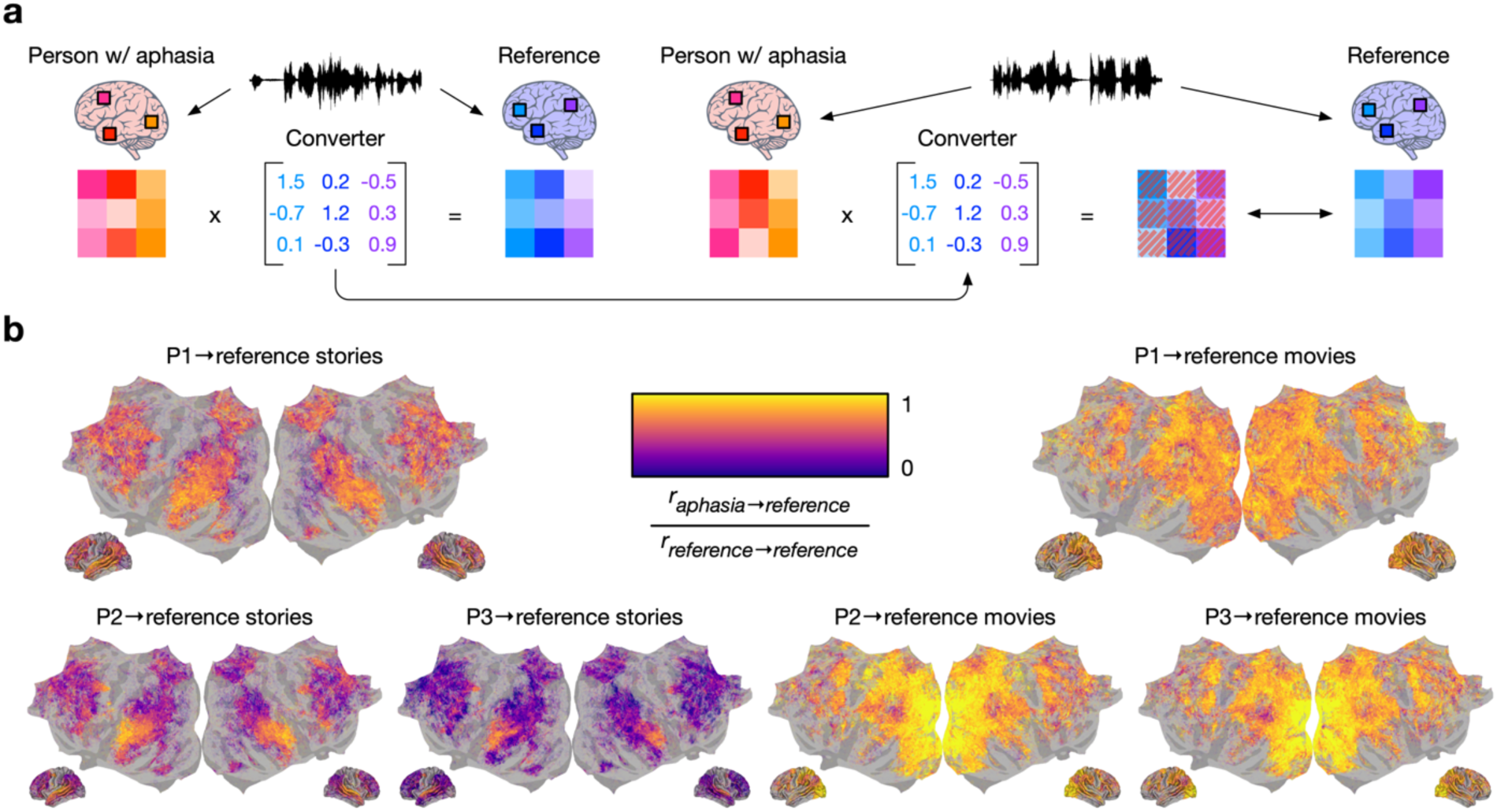
Information capacity of conceptual processing after stroke. **(a)** Cross-participant converters were trained to predict brain responses from each neurologically healthy reference participant using brain responses from each participant with aphasia. Converter performance was evaluated using the linear correlation between the predicted and actual brain responses from the reference participant. **(b)** Converters trained on narrative stories had modest performance outside of auditory and articulatory regions, indicating that some but not all of the linguistic information processed by the reference participants was also processed by the participants with aphasia. Converters trained on silent movies had high performance across cortex, indicating that most of the non-linguistic information processed by the reference participants was also processed by the participants with aphasia.

We visualized converter performance on a cortical surface atlas after averaging across reference participants^31^ (**Figure 4b**). Story-based converter performance was modest outside of auditory and articulatory regions, which indicates that some of the conceptual information in the narrative story stimuli was not fully processed by the participants with aphasia, consistent with the language comprehension impairments that we documented. Conversely, movie-based converter performance approached the ceiling in most brain regions, which indicates that most of the conceptual information in the silent movie stimuli was fully processed by the participants with aphasia. These results demonstrate that the information capacity of the semantic system can be largely spared even in participants with extensive strokes and severely impaired language processing. Consequently, semantic decoding may generalize far beyond the concepts and participants that were assessed in this study.

### Factors affecting semantic decoding performance

There are several remaining barriers to using semantic decoders as language neuroprostheses. One major barrier is that the decoders are not consistently accurate. Another major barrier is that semantic decoding from neurological patients has only been demonstrated using fMRI, which requires a large, expensive, and non-portable scanner that is unsuitable for most practical applications. To determine potential avenues for translating semantic decoders into practical language neuroprostheses, we assessed the factors that affect mental imagery decoding performance for the participants with aphasia. For these analyses, we obtained a more sensitive measure of decoding performance by assigning probabilities to the three mental imagery topics based on the decoder predictions and taking the probability of the prompted topic (**Methods**).

To make semantic decoders more accurate, we could increase the amount of training data, use higher quality brain recordings, or develop more powerful algorithms^14^. While advances in brain recording technologies and decoding algorithms are difficult to forecast, many studies have found that increasing the amount of training data consistently improves decoding performance^14,33–35^. Here we assessed whether increasing the amount of training data improves mental imagery decoding performance for the participants with aphasia (**Figure 5a**). First we found that mental imagery decoding performance increased with the amount of alignment data used to transfer decoders across participants. Notably, using silent movie data (classification probability *c* = *0.028 log_2_ m*_*movie*_ + *0.282*) was more effective than using comparable amounts of narrative story data (*c* = *0.013 log_2_ m*_*story*_ + *0.311*), demonstrating the feasibility of decoding from individuals with language comprehension impairments^16^. Next we found that mental imagery decoding performance increased with the number of reference participants (*c* = *0.034 log_2_ n*_*reference*_ + *0.414*). It is straightforward to collect large fMRI datasets from neurologically healthy participants, making this another avenue for improving decoding performance^12,36^.

**Figure 5.**
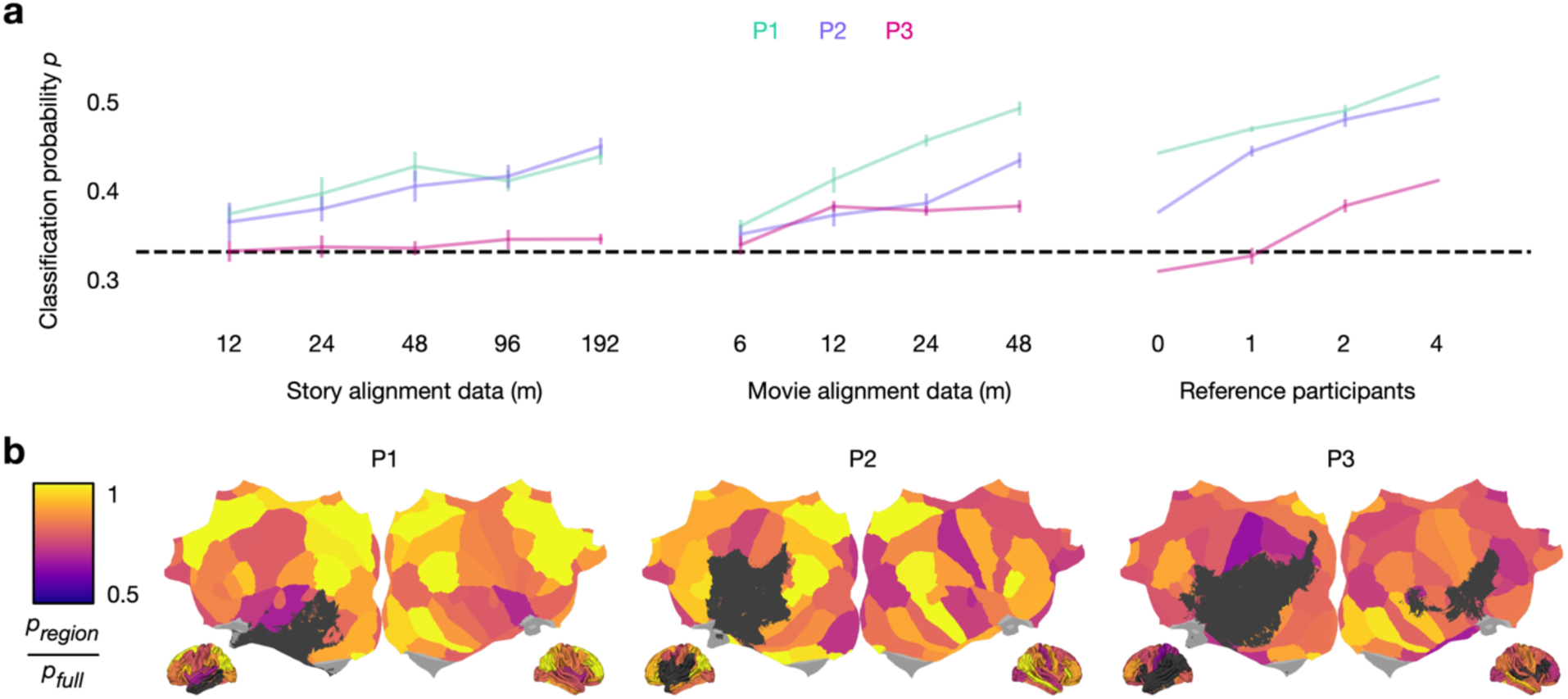
Factors affecting semantic decoding performance. **(a)** Mental imagery decoding performance increased with the amount of alignment data used to transfer semantic decoders from neurologically healthy reference participants to participants with aphasia. Aligning brain responses using silent movie data was more effective than aligning brain responses using comparable amounts of narrative story data. Mental imagery decoding performance also increased with the number of reference participants. Error bars indicate the standard error of the mean (*n* = 10 permutations). **(b)** Semantic decoders were restricted to individual brain regions to identify the potential for decoding from portable technologies that lack the full brain coverage of fMRI. Mental imagery decoding performance from specific brain regions approached decoding performance from the full brain.

To make semantic decoders more portable, we could adapt them for brain recordings made using functional near-infrared spectroscopy^37^, electroencephalography^33^, or intracranial electrodes^34,35,38^. While these more portable technologies lack the full brain coverage of fMRI, previous studies suggest that concepts are redundantly encoded in multiple brain regions^13,14^. Here we assessed whether decoding from the participants with aphasia requires the full brain coverage of fMRI. We partitioned the cerebral cortex into 68 anatomical regions and only decoded brain responses from each region (**Figure 5b**). We obtained high decoding performance from bilateral inferior parietal, medial parietal, and superior frontal regions in P1, bilateral medial parietal, right middle temporal, and bilateral inferior parietal regions in P2, and bilateral middle temporal, ventral temporal, and medial parietal regions in P3. By showing that semantic decoding does not require the full brain coverage of fMRI, these results indicate the potential for decoding from more portable technologies. By showing that the brain regions with high decoding performance can differ across individuals, these results also underscore the value of using fMRI to identify recording sites for portable decoders.

## Discussion

This study demonstrates that concept representations in individuals with post-stroke aphasia can be decoded into continuous language with a reasonable degree of accuracy. In practice, semantic decoders could be used as language neuroprostheses to predict the words, phrases, or sentences that these individuals are struggling to produce. Existing augmentative and alternative communication (AAC) devices allow individuals with aphasia to retrieve words or phrases linked to images, but they are limited in real-time responsiveness and in the scope and richness of concepts they can convey, falling short of natural communication^39^. By directly predicting what individuals are thinking about based on their brain activity, language neuroprostheses could be substantially more effective at translating rich internal thoughts into natural language.

Most existing language neuroprostheses decode motor representations to help individuals with dysarthria, who struggle to map motor representations into muscle movements^34,35,38^. However, these motor decoders are unsuitable for individuals with aphasia, who are often impaired on earlier stages of language production such as lexical access or phonological encoding. Because all stages of language production are thought to be downstream of conceptual processing, semantic decoders that target concept representations should be highly robust to the wide range of linguistic and co-occurring motoric impairments in aphasia^8–10^. Our decoding results generalized across participants with different behavioral profiles, demonstrating the potential for semantic decoding to circumvent word finding, grammatical construction, and speech motor programming impairments.

The generality of semantic decoding depends on the extent to which conceptual processing is spared in aphasia. Previous studies have approximated this extent using behavioral assessments^40–43^ and controlled neuroimaging experiments^44^, but these studies lacked the precision and coverage required to characterize the anatomical organization and information capacity of the semantic system. By using naturalistic neuroimaging experiments to measure thousands of concept representations, we found that conceptual processing can be largely spared in individuals with post-stroke aphasia despite extensive lesions impacting different regions of the semantic system. These results demonstrate that semantic decoding can be robust to the heterogeneity of lesion profiles in aphasia^3,45^.

Our results point toward several scalable avenues for improving the accuracy of the decoder predictions. It may also be possible to increase the functional utility of the decoder predictions by integrating them with residual language output, displaying them in real-time, and personalizing them for each individual with aphasia. For instance, many individuals with aphasia can produce some words despite struggling with others, so the successfully produced words could be used as linguistic context to inform the decoder predictions^46,47^. Similarly, many individuals with aphasia can successfully produce words after receiving semantic cues, so it could be helpful to display multiple sets of decoder predictions even if none are completely accurate^48^.

While this study decoded brain activity measured using fMRI, our results demonstrate that semantic decoding from individuals with aphasia does not require the full brain coverage of fMRI. Consequently, semantic decoding could be adapted for wearable technologies such as functional near-infrared spectroscopy^37^ or electroencephalography^33^. Semantic decoding using fMRI could also help plan implant locations for intracranial neuroprostheses targeting conceptual information^49,50^. Taken together, our results suggest several potential avenues for using semantic decoding to support communication in individuals with aphasia.

## Methods

### Participants

Behavioral and MRI data were collected from three right-handed participants with aphasia: P1 (male, age 55), P2 (male, age 47), and P3 (female, age 46). MRI data were also collected from four neurologically healthy participants: R1 (male, age 38), R2 (male, age 25), R3 (male, age 22), and R4 (female, age 27).

The experimental protocol was approved by the Institutional Review Board at the University of Texas at Austin. Written informed consent was obtained from all participants. Participants were compensated at a rate of $10 per hour for behavioral assessment and $25 per hour for MRI scanning. No data were excluded from the analyses.

### Behavioral assessment

In-person behavioral assessments were conducted at Austin Speech Labs and the University of Texas at Austin. Remote behavioral assessments were conducted over Zoom. All behavioral assessments were performed and scored by a trained speech-language pathologist. Speech and language processing were assessed using the extended version of the Quick Aphasia Battery^3,22^. Auditory verbal comprehension was assessed using a subset of items from the Peabody Picture Vocabulary Test^23^. Conceptual processing was assessed using the Pyramids and Palm Trees–Pictures Test^24^.

### MRI data collection

Due to a scanner upgrade at the UT Austin Biomedical Imaging Center, MRI data were collected on a 3T Siemens Skyra scanner, a 3T Siemens Vida scanner, and a 3T Siemens Prisma scanner using a 64-channel Siemens volume coil. Semantic decoders have previously been transferred across these MRI scanners^16^.

Functional data were collected using gradient echo EPI with repetition time (TR) = 2.00 s, echo time (TE) = 30.8 ms, flip angle = 71°, multi-band factor (simultaneous multi-slice) = 2, voxel size = 2.6mm x 2.6mm x 2.6mm (slice thickness = 2.6mm), matrix size = (84, 84), and field of view = 220 mm. Anatomical data for all participants except R1 were collected using a T1-weighted multi-echo MP-RAGE sequence on the same 3T Siemens Skyra, 3T Siemens Vida, and 3T Siemens Prisma scanners with voxel size = 1mm x 1mm x 1mm following the Freesurfer morphometry protocol. Anatomical data for participant R1 were collected on a 3T Siemens TIM Trio scanner at the UC Berkeley Brain Imaging Center with a 32-channel Siemens volume coil using the same sequence.

### Cortical surface reconstruction

Cortical surface meshes were generated from T1-weighted anatomical scans using Freesurfer^51^. Anatomical surface segmentations were manually checked and corrected before surface reconstruction. Functional images were aligned with the cortical surface using boundary based registration (BBR) implemented in FSL. The alignments were manually checked and corrected as necessary. Cortical surfaces were anatomically aligned with the fsAverage atlas using the *get_mri_surf2surf_matrix* function in pycortex when comparing or averaging results across participants^31,52^. Brain lesions were manually drawn in ITK-SNAP based on T1-weighted anatomical scans. Vertices corresponding to lesioned cortex were excluded from surface analyses.

### Experimental tasks

Story listening, movie watching, and mental imagery data were collected from the participants with aphasia in 5 scanning sessions. Story listening and movie watching data were collected from the neurologically healthy participants in 11 scanning sessions.

In the story listening experiment, the participants with aphasia listened to 18 narrative stories (6-16 minutes each) from *The Moth Radio Hour*. Before the experiment, the participants listened to the first 20 seconds of each story outside of the scanner. The participants were given the option to exclude stories that they struggled to understand due to speech rate, speaker accent, or audio quality. The excluded stories were replaced with other stories from *The Moth Radio Hour* chosen by the participants. The participants were also given the option to listen to the stories with subtitles. The subtitles displayed each word in black when it was spoken in the story and displayed the previous words in gray to the left of the current word. P1 and P3 chose to use the subtitles. Each story was played during a single fMRI scan with 10 seconds of silence before and after the story. The stories were played over Sensimetrics S14 in-ear piezoelectric headphones. 3 of the stories were held out for model testing (‘Where There’s Smoke’ by Jenifer Hixson, ‘Hang Time’ by Brian Gavagan, ‘The Tiniest Bouquet’ by Christine Gentry) and the remaining 15 stories were used for model training. The neurologically healthy participants listened to 54 narrative stories (6-17 minutes each) from *The Moth Radio Hour* and *Modern Love* including the stories that the participants with aphasia listened to.

In the movie watching experiment, all participants watched 13 movie clips (3-7 minutes each) from *Pixar Animation Studios* and *The Blender Foundation*. The movie clips were self-contained and almost entirely devoid of language. Each movie clip was played without sound during a single fMRI scan with 10 seconds of darkness before and after the movie clip. The original high-definition movie clips were cropped and downsampled to 727 x 409 pixels. For qualitative evaluations, 1 movie was held out for model testing (‘Sintel’) and the remaining 12 movies were used for model training. For quantitative evaluations, 3 of the movies were held out for model testing (‘La Luna’, ‘Partly Cloudy’, ‘Presto’) and the remaining 10 movies were used for model training.

In the mental imagery experiment, the participants with aphasia were prompted to silently imagine three topics—traveling to the scanning session, eating breakfast, and getting dressed. In each trial, a pseudorandom prompt was displayed for 8 seconds. The prompts indicated for the participants to imagine the corresponding topics upon seeing a thought bubble icon. The prompts also contained visual cues (a car icon for traveling to the scanning session, a plate icon for eating breakfast, and a shirt icon for getting dressed) to support comprehension. A thought bubble icon was then displayed for 30 seconds. Each topic was prompted 4 times in a single fMRI scan (7 minutes). For the mental imagery analyses, all 18 stories and all 13 movies were used for model training.

### Stimulus pre-processing

The stories and the official audio descriptions of the movies from *Pixar Animation Studios* were manually transcribed. Certain sounds (for instance, laughter and breathing) were marked to improve the accuracy of the automated alignment. The Penn Phonetics Lab Forced Aligner (P2FA) was used to automatically align the audio to the transcript^53^. Praat was used to manually check and correct each aligned transcript^54^.

### fMRI data pre-processing

Each functional run was motion-corrected using the FMRIB Linear Image Registration Tool (FLIRT) from FSL^55^. All volumes in the run were then averaged to obtain a high quality template volume. FLIRT was used to align the template volume for each run to the overall template. For participants with aphasia, the overall template was chosen to be the template for the first movie run. For neurologically healthy participants, the overall template was chosen to be the template for the first story run. The automatic alignments were manually checked.

Low-frequency voxel response drift was subtracted from the signal. Low-frequency voxel response drift in the story and mental imagery scans was identified using a 2nd order Savitsky-Golay filter with a 120 second window. Because the movie scans were shorter, low-frequency voxel response drift was identified using a Legendre polynomial of degree three^56^. The remaining signal was then *z*-scored.

### Language model

GPT (also known as GPT-1) is a 12-layer transformer neural network trained to predict the next word in a sequence given the previous words^57^. In order to perform this task, GPT learns to encode the meanings of input sequences in its hidden layers. Given a word sequence *S* = (*s_1_*, *s_2_*, … , *s*_*n*_), the output from GPT was used to model the probability distribution over the next word *s*_*n*+_*_1_* and the hidden layers of GPT were used to model the meanings of the previous words (*s_1_*, *s_2_*, … , *s*_*n*_).

### Encoding model

Voxelwise encoding models were trained to predict brain responses given word sequences. Quantitative features were extracted for each word-time pair (*s*_*i*_, *t*_*i*_) by providing the word sequence (*s*_*i*–_*_5_*, *s*_*i*–_*_4_*, … , *s*_*i*–_*_1_*, *s*_*i*_) to the GPT language model and taking the activations from the ninth hidden layer^25,26,58,59^. This yields a new list of vector-time pairs (*M*_*i*_, *t*_*i*_) where *M*_*i*_ is a 768-dimensional embedding for *s*_*i*_. These vectors were then resampled at times corresponding to the fMRI acquisitions using a three-lobe Lanczos filter.

A linearized finite impulse response (FIR) model was fit to every cortical voxel in the participant’s brain. A separate linear temporal filter with four delays (*t* − *1*, *t* − *2*, *t* − *3*, and *t* − *4* timepoints) was fit for each of the 768 features, yielding a total of 3,072 features. With a TR of 2 seconds this was accomplished by concatenating the feature vectors from 2, 4, 6, and 8 seconds earlier to predict responses at time *t*. Each feature channel of the training matrix was *z*-scored before the regression procedure.

The 3,072 weights for each voxel were estimated using L2-regularized linear regression. The regression procedure has a single free parameter that controls the degree of regularization. This regularization coefficient was found by repeating a regression and cross-validation procedure 50 times. In each iteration, approximately a fifth of the timepoints were removed from the model training dataset and reserved for validation. Then the model weights were estimated on the remaining timepoints for each of 10 possible regularization coefficients (log spaced between 10 and 1,000). These weights were used to predict brain responses for the reserved timepoints, and *R*^2^ was computed between the predicted and actual brain responses. The regularization coefficient was chosen as the value that led to the best performance, averaged across bootstraps, voxels, and neurologically healthy participants, on the reserved timepoints. After training, the encoding model can take any word sequence *S* and predict brain responses *R*^4^(*S*).

### Cross-participant transfer

To reduce the amount of data required from the participants with aphasia, encoding models were trained on brain responses from the neurologically healthy reference participants and then transferred to the participants with aphasia. Because the anatomical structure and functional organization of the brain vary across individuals, transferring models across participants requires aligning their brain responses^14,60^.

To learn this alignment, brain responses *R*_*reference*_ and *R*_*ap*_*_ℎ_*_*asia*_ were collected while each reference participant and participant with aphasia were shown a common set of story and movie stimuli. A cross-participant converter *C*_*reference* → *ap*_*_ℎ_*_*asia*_ was trained to predict the activity in each voxel of the participant with aphasia using the activity in the 10,000 best predicted voxels of the reference participant. The converter weights were estimated using L2-regularized linear regression. The regularization coefficient was found by repeating the regression and cross-validation procedure 50 times for each pair of reference participants. The regularization coefficient was chosen as the value that led to the best performance, averaged across bootstraps, voxels, and reference participant pairs, on the reserved timepoints. After training, the converter can take any encoding model trained on the reference participant and align it with brain responses from the participant with aphasia.

Converters were also used to identify concept-selective voxels in the participants with aphasia. Converters were trained on brain responses to either stories or movies and then evaluated by aligning brain responses to the other modality. Prediction performance was quantified using the linear correlation between the aligned and actual brain responses from the participant with aphasia. Prediction performance was *z*-scored and averaged across modalities to identify the 10,000 voxels with the highest cross-modality prediction performance.

For some analyses, separate converters *C*_*ap*_*_ℎ_*_*asia* → *reference*_ were trained to predict the activity in each voxel of the reference participant using the activity in the concept-selective voxels of the participant with aphasia. Converter performance was used to quantify whether the information processed in each brain region of the reference participant was also processed somewhere in the participant with aphasia.

### Word rate decoder

A word rate decoder was estimated for each participant with aphasia to predict when concepts were perceived or imagined. The word rate at each fMRI acquisition was defined as the number of stimulus words that occurred since the previous acquisition.

To identify the voxels that encode word rate, L2-regularized linear regression was used to estimate a set of weights that predict brain responses *R* from word rate *w*. A linear temporal filter with four delays (*t* − *1*, *t* − *2*, *t* − *3*, and *t* − *4* timepoints) was fit for the word rate feature. With a TR of 2 seconds, this was accomplished by concatenating the word rate from 2, 4, 6, and 8 seconds earlier to predict the brain responses at time *t*. The 3,000 voxels with the highest cross-validation performance were selected for word rate decoding.

To train the word rate decoder, L2-regularized linear regression was used to estimate a set of weights that predict word rate *w* from brain responses *R*. A separate linear temporal filter with four delays (*t* + *1*, *t* + *2*, *t* + *3*, and *t* + *4* timepoints) was fit for each selected voxel. With a TR of 2 seconds, this was accomplished by concatenating the brain responses from 2, 4, 6, and 8 seconds later to predict the word rate at time *t*. Given new brain responses *R*_*ap*_*_ℎ_*_*asia*_, the model predicts the word rate at each acquisition. The time between consecutive acquisitions (2 seconds) can then be divided by the predicted word rates (rounded to the nearest non-negative integers) to predict word times.

### Semantic decoder

The goal of language decoding is to maximize the probability distribution *P*(*S* | *R*) over word sequences *S* given brain responses *R*. *P*(*S* | *R*) can be factorized into the product of a prior distribution *P*(*S*) over word sequences and an encoding distribution *P*(*R* | *S*) over brain responses given word sequences^14,61^.

*P*(*S*) was estimated using the language model. The language model maps from a word sequence to the probability distribution over the next word. The probability of observing *S* = (*s_1_*, *s_2_*, … , *s*_*n*_) in natural language can be modeled by multiplying the probabilities *P*(*s*_*i*_ | *s_1_*_:*i*–*1*_) of each word conditioned on the previous words.

*P*(*R* | *S*) was estimated using the encoding models. Each encoding model maps from semantic features of *S* to brain responses *R*^4^(*S*). Assuming that BOLD signals are affected by Gaussian additive noise, *P*(*R* | *S*) can be modeled as a multivariate Gaussian distribution with mean *μ* = *R*^4^(*S*) and covariance ∑ = ⟨(*R* − *R*^4^(*S*))^*T*^(*R* − *R*^4^(*S*))⟩^62^. Because the aligned encoding models from the neurologically healthy reference participants could only explain a small fraction of the variance in the brain responses from the participant with aphasia, ∑ was approximated by computing *R*^*T*^*R* for each training story from the participant with aphasia and averaging the brain response covariance matrices across the training stories.

To decode new brain responses *R*_*ap*_*_ℎ_*_*asia*_, the beam search decoding approach introduced in ref.^14^ was used to generate word sequences with high values of *P*(*S*) and *P*(*R*_*ap*_*_ℎ_*_*asia*_ | *S*). The decoder maintained a beam containing the *k* most likely word sequences up to each timepoint. Upon receiving new brain images, the word rate model was used to detect new words. Next the language model was used to generate continuations for each candidate *S* in the beam by predicting the probability distribution *P*(*s*_*n*_ | *s*_*n*–*i*_, … , *s*_*n*–_*_1_*) over the next word *s*_*n*_ given the words predicted in the previous 8 seconds (*s*_*n*–*i*_, … , *s*_*n*–_*_1_*). Each word in the language model nucleus^63^ was appended to the candidate to form a continuation *H*. Finally the aligned encoding models from the reference participants were used to rank the continuations based on the likelihood *P*(*R*_*ap*_*_ℎ_*_*asia*_ | *H*) of observing the brain responses. The *k* most likely continuations were retained in the beam for the next timepoint and the process was repeated until the entire scan was decoded.

Previous studies have found that combining models from multiple reference participants can improve decoding performance^16,60^. As a result, *P*(*R*_*ap*_*_ℎ_*_*asia*_ | *H*) was separately computed using the aligned encoding model from each reference participant and the ranks were averaged. When decoding brain responses to stories and movies, the likelihoods *P*(*R*_*ap*_*_ℎ_*_*asia*_ | *H*) were also averaged with the likelihoods *P*(*R*_*ap*_*_ℎ_*_*asia* → *reference*_ | *H*), which were obtained by multiplying the brain responses *R*_*ap*_*_ℎ_*_*asia*_ by the cross-participant converters *C*_*ap*_*_ℎ_*_*asia* → *reference*_ and evaluating the aligned responses using the native encoding model from each reference participant^16^. This regularization leverages the fact that the noise covariance ∑ can be estimated using many more hours of brain responses from the reference participants. When assessing the benefits of cross-participant modeling, aligned encoding models from the reference participants were compared to native encoding models from the participant with aphasia. Custom analysis software was written in Python using the NumPy^64^, SciPy^65^, PyTorch^66^, Transformers^67^, and pycortex^52^ libraries.

### Decoding performance

Decoder predictions from the story and movie experiments were compared to target words from the stimulus transcripts using the BERTScore metric^28^. BERTScore was computed between the predicted and target words within a 20 second window around each second of the stimulus. BERTScore uses a bidirectional transformer language model to represent the meaning of each predicted and target word as a contextual embedding, and scores each target word based on its maximum cosine similarity across all of the predicted words. These scores are averaged across the target words to quantify the similarity of meaning between the predicted and target word sequences.

Because BERTScore values are not inherently interpretable, they were normalized with respect to a null distribution^28^. Null sequences were generated by sampling from the language model without using any brain data except to predict word times^14^. The null model maintained a beam of 10 candidate sequences and generated continuations from the language model nucleus at each predicted word time^63^. The only difference between the actual decoder and the null model was that, instead of ranking the continuations by the likelihood of the fMRI data, the null model randomly assigned a likelihood to each continuation. For each participant with aphasia, this process was repeated 200 times to generate 200 null sequences. This process was as similar as possible to the actual decoder without using any brain data to select words, so these sequences reflected the null hypothesis that the decoder does not recover meaningful information about the stimulus from the brain data. The null sequences were scored against the target words to produce a null distribution of decoder scores for each participant with aphasia.

Decoder predictions from the mental imagery experiment were compared to the prompted topics (“I traveled here”, “I got dressed”, and “I ate breakfast”) using BERTScore. Because the participants could have imagined idiosyncratic concepts for each topic, they were also instructed to describe what they imagined after the scan. P1 and P2 described what they imagined out loud while P3 wrote what she imagined. The descriptions were manually edited to remove filler words and define proper nouns, and were then concatenated with the prompts to form target words for each topic. To evaluate accuracy, BERTScore was computed between the decoder predictions and the target words for the prompted topic. To evaluate discriminability, BERTScore was computed between the decoder predictions and the target words for all three topics, and the similarities were normalized into probabilities that the participant was imagining each topic. To make the decoding results more stable, the decoding procedure was repeated for 5 sets of 10,000 voxels sampled from the top 12,500 concept-selective voxels and the probabilities were averaged across the samples.

### Anatomical regions

Whole brain MRI data were partitioned into 4 anatomical regions using Freesurfer labels: anterior left hemisphere, posterior left hemisphere, anterior right hemisphere, and posterior right hemisphere. Anterior regions were defined using the *superiorfrontal*, *rostralmiddlefrontal*, *caudalmiddlefrontal*, *parsopercularis*, *parstriangularis*, *parsorbitalis*, *lateralorbitofrontal*, *medialorbitofrontal*, *precentral*, *paracentral*, *frontalpole*, *rostralanteriorcingulate*, and *caudalanteriorcingulate* labels. Posterior regions were defined using the *superiorparietal*, *inferiorparietal*, *supramarginal*, *postcentral*, *precuneus*, *posteriorcingulate, isthmuscingulate, superiortemporal*, *middletemporal*, *inferiortemporal*, *bankssts*, *fusiform*, *transversetemporal*, *entorhinal*, *temporalpole*, *parahippocampal*, *lateraloccipital*, *lingual*, *cuneus*, and *pericalcarine* labels. Whole brain MRI data were also partitioned into 34 anatomical regions in each hemisphere using these labels along with the *insula* label in order to identify brain regions with high mental imagery decoding performance.

### Functional networks

Whole brain MRI data were partitioned into 5 functional networks using a clustering approach^26^. Encoding models were trained to predict brain responses from the neurologically healthy participants using lexical semantic features of the stimulus words^13^. Encoding model weights were averaged across the 4 delays to produce a conceptual tuning vector for each voxel, which captures how the voxel responds to the stimulus features. The top 10,000 voxels were identified in each reference participant based on cross-validation performance. The conceptual tuning vectors of the top 10,000 voxels were concatenated across participants and rescaled to have unit norm. Principal components analysis was applied to the conceptual tuning vectors, and 64 dimensions that explained 80% of the variance were chosen.

Spherical *k*-means clustering was applied to the principal components. To determine the number of clusters, the inertia of the clustering algorithm was computed for a range of clusters between 1 and 20. This was used to identify the point where the inertia changes from an exponential drop to a linear drop in inertia. This point occurred at 5 clusters. Each of the 5 clusters was interpreted by projecting the lexical semantic features of 10,470 words onto the cluster centroid. The clusters were subjectively determined to represent *concrete*, *social*, *place*, *temporal*, and *people* concepts.

The conceptual tuning vectors were anatomically aligned with the fsAverage atlas and averaged across neurologically healthy participants. The conceptual tuning vectors were then projected onto the 64 principal components and then assigned to one of the 5 clusters. This process results in a functional network label for every fsAverage vertex.

### Statistical testing

Statistical significance of decoder scores was assessed using a one-sided non-parametric test. The decoder scores were averaged across the stimulus windows and the observed mean decoder score was compared to the mean decoder scores for the null sequences. *p* values were computed as the fraction of null sequences with a mean decoder score greater than or equal to than the observed mean decoder score.

Statistical significance of encoding model performance was assessed using a one-sided permutation test. The encoding models were used to predict brain responses to 3 held-out test stories. Prediction performance for each voxel was computed as the linear correlation between the predicted and actual response time-courses. A null distribution for each voxel was obtained by randomly resampling (with replacement) 10-TR blocks from the voxel’s actual response time-course. Resampling contiguous blocks preserves the auto-correlation structure of the voxel’s responses. The null prediction performance was then computed as the linear correlation between the predicted and permuted response time-courses. Repeating this process for 1,000 trials provided a null distribution of prediction performance for each voxel. *p* values were computed for each voxel as the fraction of permutations with prediction performance greater than or equal to than the observed prediction performance.

Statistical significance of tuning correlation was assessed using a one-sided permutation test. To compare conceptual tuning between a participant with aphasia and the neurologically healthy participants, lexical semantic encoding models weights were averaged across the 4 delays to produce a conceptual tuning vector for each voxel. The conceptual tuning vectors were anatomically aligned with the fsAverage atlas. The lexical semantic features of 10,470 words were projected onto the conceptual tuning vectors from the participant with aphasia to simulate how each cortical location would respond. This process was repeated for the average conceptual tuning vectors across the neurologically healthy participants. Tuning correlation was computed by taking the rank correlation between the two sets of simulated responses for each fsAverage vertex and then averaging across fsAverage vertices. A null distribution was obtained by permuting the conceptual tuning vectors across fsAverage vertices before simulating brain responses. Repeating this process for 1,000 trials provided a null distribution of tuning correlation. *p* values were computed as the fraction of permutations with a tuning correlation greater than or equal to than the observed tuning correlation.

Quantitative tuning correlation analyses were restricted to concept-selective vertices in the fsAverage atlas. These vertices were identified by training a lexical semantic encoding model for each neurologically healthy participant. Concept-selective voxels were identified as voxels with an observed prediction performance score that was significantly higher than its null distribution. The binarized voxel masks of conceptual selectivity were anatomically aligned with the fsAverage space and averaged across neurologically healthy participants. Vertices with a mean value greater than 0.5 were considered to be significantly predicted, resulting in 74,370 significantly predicted vertices out of 327,684 total vertices.

Unless otherwise stated, all tests were performed within each participant and then replicated across all participants (*n* = 3). All tests were corrected for multiple comparisons when necessary using false discovery rate^68^.

## Data availability

All data used in the analyses will be made publicly available at OpenNeuro before publication.

## Code availability

All code used in the analyses will be made publicly available at GitHub before publication.

## Acknowledgements

We thank the participants and their loved ones for their time, dedication, and generosity.

## Author contributions

Conceptualization: J.T., A.G.H., M.L.H.; Methodology: J.T., S.S., S.M.W., A.G.H., M.L.H.; Software and resources: J.T.; Investigation and data curation: J.T., C.M., A.C., L.D.W., and J.A.; Formal analysis and visualization: J.T.; Writing (original draft): J.T.; Writing (review and editing): J.T., C.M., A.C., L.D.W., J.A., S.S., S.M.W., A.G.H., M.L.H.; Supervision: A.G.H., M.L.H.

## Materials & Correspondence

Correspondence and requests for materials should be addressed to Jerry Tang, Alexander G. Huth, and Maya L. Henry.

## Supplementary Information

**Supplementary Table 1.** Decoder predictions for a perceived story. Semantic decoders were evaluated on single-trial BOLD fMRI responses recorded while three participants with aphasia listened to the test story ‘Where There’s Smoke’ by Jenifer Hixson from *The Moth Radio Hour*. The actual stimulus words are shown alongside the decoder predictions for each participant.

|  | Actual stimulus | P1 | P2 | P3 |
| --- | --- | --- | --- | --- |
| 0 | i reached over and secretly undid my seatbelt and when his foot hit the brake at the red light i flung open the door and i ran |  |  |  |
| 10 | i had no shoes on i was crying i had no wallet but i was ok because i had my cigarettes | she said i should tell my wife to get a divorce she was crying so much it wasn't funny it was a joke i laughed like | i am an engineer for the company that was founded by a woman i met online who is the author of a novel i think was in the | we are very close in age i was born into the us and went to college there were three schools of study abroad one was a state |
| 20 | and i didn't want any part of freedom if i didn't have my cigarettes when you live with someone who has a temper a very bad temper a | a little girl i was just being silly but the laughter was genuine so i started to get excited it was all about fun i had always been good at | us originally it was a comedy series and was supposed to be the first book the film was not a very popular movie the producers said it | university i didn't attend they taught us all that we could and we learned we were a bit of a minority they gave us a chance to |
| 30 | very very bad temper you learn to play around that you learn this time i'll play possum and next time i'll | laughing and being funny in high school my best friend was a guy from a bad family he was the oldest of four brothers and two | was a comedy the actors didn't agree with it but i think the plot is a good one we all know the ending is funny and it makes sense the other | grow a few more seeds and we did we grew it into a tree that looked like it was made of glass the fruit was green but not purple |
| 40 | just be real nice or i'll say yes to everything or you make yourself scarce or you run and this was one of the times when you just run and as | sisters i was his best friend in high school we grew apart from each other over years we went on walks and i helped her | way around we have a different perspective on life in general the universe has changed from what it was the other day and i | or orange in color and was perfectly normal looking for someone who is really into that kind of thing but i was |
| 50 | i was running i thought this was a great place to jump out because there were big lawns and there were cul de sacs and sometimes he would come after me and drive and yell stuff at me to get back in get back in | get to the top of the cliff i made her climb a hundred feet into the air before landing on the rocks and rolling back down to the | have decided to stop living in the past i know my family has moved away from home and that they are | always so worried about people going out on their own and not having friends who were into girls or even guys but i never considered myself a bad |
| 60 | and i was like no i'm out of here this is great and i went and hid behind a cabana and he left and i had my cigarettes and uh i started to walk in this | bottom again if you can't catch yourself or the person below you they will die the same as anyone else i saw my parents do so that we could learn about ourselves and | still living on their own there is nothing worse than being homeless you know how to survive in the dark i mean seriously that's like the best you could come up | person because i had a hard time loving people who were mean to me or just treated me poorly i started seeing my dad more frequently he |
| 70 | beautiful neighborhood it was ten thirty at night and it was silent and lovely and there was no sound except | have the experience of our youth to share with us the lessons they had taught us on this journey we could see it all and | with right there so yeah i'm pretty much at a dead end and i had to go through all that i was living on | had a new car but didn't use it anymore he got better and the same old thing happened again the only difference is he wasn't living on the streets |
| 80 | for sprinklers ch ch ch ch ch ch ch and i was enjoying myself | it was like a dream so i just fell asleep it felt really weird as | in my own room and had to take a shower so i got a bunch of dirty clothes and | or a cheap hotel room like the other homeless people so i don't know i get home and i ask my landlord |
| 90 | and enjoying the absence of anger and enjoying these few hours i knew i'd have of freedom and just to perfect it i thought i'll have a smoke and | if my brain just blew itself out | my pants were ripped up the back of her head | for the rent check to make sure i didn't get it wrong then |
| 100 | then it occurred to me with horrifying speed i don't have a light just then as if in answer i see a figure up ahead | and it wasn't like it had been burned off so i don't know how it got in there i couldn't find it the next day when i came back and looked up | with a sharp edge that makes the blood flow faster than the brain is designed to take it's place so now we can do our research but in the process | he says he doesn't know about it because it wasn't in the house so it was the next day when i got to the door i |
| 110 | who is that it's not him ok they don't have a dog who is that what uh what are they doing out on this suburban street and the person comes closer and i could see it's a woman | the owner was gone my friend who lived down the street from me was at work when we saw a police officer | of the experiment i am thinking about the fact that we are not yet able to understand how we can help them get their | was at the time i had already decided on the name and date of my engagement party for the |
| 120 | and then i can see she has her hands in her face oh she's crying and then she sees me and she composes herself and she gets closer and i see she has no | in our neighborhood walking up to a girl on the corner with her boyfriend a friend of mine came over and told them that their girlfriend was waiting for them she asked | lives back on track after all of this my family and i were so afraid to see what was out there that we would spend every day on our own | month we got together i asked him to give me a hug and a kiss after we broke apart he said i can only say one thing in that |
| 130 | shoes on she has no shoes on and she's crying and she's out on the street street i recognize her though i've never met her and just as she passes me she | us to tell her about the guy we knew from high school who was really cute and i said that was not the same person but i did know him he asked if i | at home we were still learning how to get around so we could work together but it was hard when you are not paying rent and your car | case i thought i could talk to you about the matter with a fellow officer of the court it was a long process the prosecutor and i were both |
| 140 | says you got a cigarette and i say you got a light and she says damn i hope so and then sh first she digs into her cutoffs in the | was his girlfriend i told him no i don't have one and he said well i will do it then he says you better give it to me i | is broke i would ask if she was homeless she said no i don't care you can borrow mine i told her she didn't have | required to participate in the questioning which i was told was necessary as a witness the officer said we can see you now sir the guy |
| 150 | front nothing and then digs in the back and then she has this vest on that has fifty million little pockets on it and she's checking and checking and it's looking bad it's looking very bad | took it out and put it in a plastic bag with a piece of duct tape to wrap it around then | one i looked at my wallet and saw no credit card money or change in it it was all a bit strange so | behind us says there are people coming this way he knows that his friends are waiting on him and he runs off with them |
| 160 | she digs back in the front again deep deep and she pulls out a pack of matches that had been laundered at least once ukgh we open | i grabbed a metal pipe and cut it into about inches thick with a large length of steel tubing in the top half i put the metal | i was walking home and noticed my bike had been replaced by a car with a huge tire iron sitting on the hood i pulled up | so i just let them get away i mean if you're going to make them feel bad for leaving you out on a date and then be left alone |
| 170 | it up and there is one match inside ok oh my god this takes on it's like nasa now we got to like oh how are we gonna do it ok and we we hunker down | rod on the bottom part and then cut it into small pieces i have a ton of pain | the back of the truck and grabbed the gun off the passenger seat as we were rolling down the hill | in the restaurant when they came to pay i got a call from a local police department that was on a busy |
| 180 | we crouch on the ground and where's the wind coming from we're stopping i take out my cigarettes let's get the cigarettes ready oh my brand she says not surprising and | killers and am constantly tripping up or coughing in the heat of a room so the police just let it go they tried to force | to the driveway i hit the brakes and started honking my horn at her she immediately responded by pointing the car in front of her to | street outside of a major shopping center with an old fashioned subway system in the background there was a train station that i remember the conductor |
| 190 | we both have our cigarettes at the ready she strikes once nothing she strikes again yes fire puff inhale mm sweet kiss of that cigarette | the door shut and he pushed it back a few inches then shoved my foot down hard on the knob so i slammed | move it back a little bit and then slowly pull it forward a few inches again she says no please just take a breath she leans down | had a big box of popcorn and a huge plastic cup filled with a bright red juice i was in a daze the |
| 200 | and we sit there and we're loving the nicotine and we both need this right now i can tell the night's been tough immediately we start to reminisce | the door and started pushing it back open i was feeling pretty bad after that i walked into my apartment to see a stranger | to kiss me on the lips and i feel this intense moment that i can't describe as a love or a passion i want to say i | whole time and was trying to process it all i realized i didn't feel anything so i took off the shoes |
| 210 | about our thirty second relationship i didn't think that was gonna happen me neither oh man that was close oh i'm so lucky i saw you yeah then she | on the phone who had just arrived in town the woman said hi i live here too and i was like hey are | have a strong feeling that you should not be alone i know you don't need anyone else to do this but you are and | and my socks i don't think anyone would ever find them but i guess i didn't care so i pulled on the shirt and |
| 220 | surprises me by saying what was the fight about and i say wha what are they all about and she said i know what you mean she said was it a bad one and and i said | you the girl that came over i asked yeah is this your girlfriend i said i don't know why you think so | i can feel you looking at me i'm not talking about me it's something that happened when i was around a lot of people had similar experiences | it came out just like i said it did but now i don't see the hole or anything else the guy is saying it |
| 230 | you know like medium she said oh and we start to trade stories about our lives we're both from up north we're both kind of newish to | it's a very basic concept in our society where we have no knowledge of the world outside the us there are many countries with | in their life it was like living on the edge of an infinite cliff that dropped into a very dark and | doesn't matter what happens next but i just know i will never see her again the thought of her being at a nursing home for years in |
| 240 | the neighborhood this is in florida we both went to college not great colleges but man we graduated and i'm actually finding myself a little jealous of her because she has this really cool | large cities and towns in the united states where i lived that had a military base but only one major airport which was not used for | terrifying valley below me a young man i knew at first saw a large open field he had a big wooden crate of beer | an insane asylum or a hospital with a mental health institution and a psychiatric ward my dad has a huge medical insurance policy on everything |
| 250 | job washing dogs she had horses back home and she really loves animals and she wants to be a vet and i'm like man you're halfway there | business or anything else he didn't seem like the type to take orders he just kept his head down and would have been happy with a sandwich but then he | and some food in his lap i told him it was too dangerous i didn't want to leave him alone with it i said | including his retirement pension i would recommend a large house with a swimming pool for guests to enjoy on weekends and vacations like the one at |
| 260 | i'm a waitress at an ice cream parlor so um that's not i don't know where i want to be but i know it's not that and then it gets a little deeper | said no you cannot i can't i love my job and i can help with anything i want i | i don't know i told him no i didn't have a dog so i could keep him around i was able to make | your hotel i have a room service cart for the two of you and i will make sure you get it you both have |
| 270 | and we share some other stuff about what our lives are like things that i can't ever tell people at home this girl i can tell her the really ugly stuff and she | could teach a class or a group of friends a story and a few words i had heard at some point the | myself more comfortable as i moved forward with the baby and he was fine but the feeling of being trapped inside his body | to take turns telling us what you need for your wedding dress and then getting it made i told him it would probably cost me |
| 280 | still understands how it can still be pretty she understands like how nice he's gonna be when i get home and how sweet that'll be | night before i woke up to a note on the bed i turned around and saw her face in the mirror i told her what had happened i said | was still very much present as he lay in bed i noticed how he looked and felt every part of him was now all covered with his clothes i said to | months to make but it wasn't worth that i could be happy for him and just not care about myself or his life i was really |
| 290 | we are chain smoking off each other oh that's almost out come on and we go through this entire pack until it's gone and then i say you know what uh this is a little funny but you're gonna | to her she just shook her head i took my finger and pushed it in her mouth she pulled her hand away | him are you ok my dad replied just fine he asked me how much for a pair of shorts i told him | glad when my mom moved away from me because i wasn't ready for the situation at first i was in a very |
| 300 | have to show me the way to get home because although i'm twenty three years old i don't have my driver's license yet and i just jumped out right when i needed to and she says well why don't you come back to my house and i'll give you | i let it fall to my side and walked out i got about feet down the road before the car pulled in front of me i jumped back up | to put his money back in my wallet he said i wouldn't pay him for it because he had stolen it and that he didn't want it | strange place i had just moved into the house i didn't know anyone was there at this point but i figured if i went inside it would |
| 310 | a ride i say ok great and we start walking and uh we get to this um lots of uh lights and uh the roads are getting | on the porch and yelled hey you guys can come out so we all climbed down i said ok let me check i walked through the woods to the top of the hill | back but he did it anyway so i thought well here we go this was one of those things i remember i was driving down a major highway that is | be ok because i couldn't see anything and then i realized i had left a note in my purse but it was gone i thought that was it until i |
| 320 | wider and wider and there's more cars and i see um lots of stores you know laundromats and dollar stores and emergenc centers and | and the forest opened out into a massive valley filled with huge buildings in various stages of construction and | lined with small towns on either side of a mountain range in the south or the us the air force | looked around and saw a car pulling into a gas station that was shut down by a |
| 330 | then we cross over us one and uh she leads me to some place and i think no but yes carl's efficiency | structures like a giant pyramid or an ancient city on the planet earth but it's not that simple i do know of a | and other military will be coming from both sides to protect you the people of the us need you in the | police officer this happened at a local high school in a neighborhood near a college town that has a football stadium |
| 340 | apartments this girl lives there and it's horrible and it's lit up so bright just to illuminate the horribleness of it it's the kind of place where you drive your car right up and the door's | guy who lived in the us for years before moving here from a different country that was just starting | middle i don't think there are very many jobs available for this level of work in the world that | a bowling alley a golf course a tennis court a soccer field a ski hill a small community college |
| 350 | right there and there's fifty million cigarette butts outside and there's like doors one through seven and you know behind every single door there's some horrible misery going on there's someone crying or drunk | out to build on the same principles and values of the two systems that made up the human race as an entire being a god with no soul | require more time to think and process it all i have been at a place where i could spend hours in the library or my apartment i can walk up to | with a school bus that runs a ton of people over a few hours but for a year and a half or more you can make |
| 360 | or lonely or cruel and i think oh god she lives here how awful we go to the door door number four and she very very quietly | or purpose but a creator all the same we are a machine and a tool of evil so it is a simple task to make a device that will work but can only | the top and get out of the way but it's too high i need a ladder so i start climbing down it slowly and then i feel | the time you have left to enjoy the things you are experiencing as you go through the process |
| 370 | keys in as soon as the door opens i hear the blare of television come out and on the blue light of the television the smoke of a hundred cigarettes in that little crack of light | function on a single component and which requires the input of the system to generate the data for the application process | this sensation as i reach the top my mind is flooded with a huge amount of images and emotions that i couldn't even comprehend for | of making the changes i would also suggest you write up a check for a year to get your wife and the kids a free vacation so |
| 380 | and i hear the man and he says where were you and she says never mind i'm back and he says you alright and she says yeah i'm alright | and then he said i don't know you so i asked him who is he i told him it's me he says | the time we were there he asked if i would be ok i said no but he kept saying to me is your girlfriend okay i | they can spend their time together we decided that we would be happy with just her and me but the girls wanted to come along so we did we started talking |
| 390 | and then she turns to me and says you want a beer and he says who the fuck is that and she pulls me over and he sees me and he says oh hey i'm not a threat | no i'm sorry and turns around i tell him he better leave now he goes i told him to go he looks at me and | replied he looked at me like i'm stupid he said no i told him to shut up he didn't and he took out a | about how much we enjoyed playing video games together she was pretty cute and we really had fun but i just didn't want to make things weird so we started |
| 400 | just then he takes a drag of his cigarette a very hard hard drag you know the kind that makes the end of it really heat up hot hot and long and it's a little scary and | says no sir he then takes his little girl with him he tells her to put her arm around my neck | pair of handcuffs i think they were a size or two thick the cuffs were wrapped in duct tape and a large belt loop around the ankles | talking a lot she was super excited about her boyfriend and her life was amazing she would ask me questions like when does it stop hurting what is the cause of |
| 410 | i follow the cigarette down because i'm afraid of that head falling off and i'm surprised when i see in the crook of his arm a little boy sleeping | and walk down the sidewalk in the middle of a busy road that leads to an outdoor bar or some other venue for the evening | that held them tight to the ground this would prevent them from sliding away as they did in my experience with these things so they could escape into the | pain and how did you deal with it all those questions i had to ask about this thing that was in the water i thought it was a fish |
| 420 | a toddler and i think and just then the girl reaches underneath the bed and takes out a carton and she taps out the last s pack of cigarettes in there | the night before a huge gathering with a large table set up and a giant buffet style meal of food | forest i went to a place i knew and we spent the next months doing all the things we had planned out in the | and i didn't even notice the hole i had dug in the side of my car that was leaking |
| 430 | and on the way up she kisses the little boy and then she kisses the man and the man says again you alright and she says yeah i'm just gonna go out and smoke with her and so | spread out all around the table with everyone enjoying it and the only thing that had me thinking was how awesome this whole experience is but it | last year of college and the day was finally here i was walking along a quiet street looking | water into the hole i couldn't see where it came out i didn't really feel the pain in the pit of |
| 440 | we go outside and sit amongst the cigarette butts and smoke and i say wow that's your little boy and she says yeah isn't he beautiful and i say yeah he is he is beautiful | also makes me wonder what is so special about the other two things that you said to me i asked you how old is your son | at houses in a park and a young boy approached me i said no thanks i don't know you but i can't walk very | my stomach that was causing it at the time but i was still angry with the girl and my dad was |
| 450 | he's my light he keeps me going she says we finish our cigarettes she finishes her beer i don't have a beer cause i can't go home | he says i am not his father but i think he means you are my daughter he is going to marry you in six months if | far right now i'm going to sleep for a bit so i can make a plan i do a little more than an hour | never a fan of mine so i went to a local college where we had a large library with about books |
| 460 | with beer on my breath and she goes inside to get the keys she takes too long in there getting the keys and i | not seven then eight and when they get back to the house i run out the door | or two and then we got a call that someone was getting away from us the driver told | and tons of free time available so the next day i go into work to make sure i have a good night and |
| 470 | think something must be wrong and she comes out and she says look i'm really sorry but um like we don't have any gas in the car it's already on e and he needs to get to work in the morning and um | and look back for her car keys she didn't get them from the counter i | me i should just stop because they didn't want me to drive them home so they can get some food they | it was the perfect opportunity i was ready for a weekend trip to a country club or some such but when we |
| 480 | i you know i i'm gonna be walk to work as it is so what i did was though here look i drew out this map for you and you're really you're like a mile and a half from home and um if you walk three streets over you'll be back on that pretty street | ask if she needs a ride i said she should just park on the street i can go up the hill to the front of the building and back down | say i should have the truck i said no they insist i take the car they want the keys to the house and the money back in the glove | got there i realized that we were being filmed in a different area so we could see the people from the airport who were |
| 490 | and you just take that and you'll be fine and she also has wrapped up in toilet paper seven cigarettes for me a third of her pack i note and a new pack of | the other side so the alley is blocked off with a few large garbage cans i grab a roll of duct tape and put it to her mouth while i get the | box because they can put it on their credit card that is just like selling a piece of jewelry with a fake diamond and the same goes for the ring it was a wedding | about to leave we then saw that a guy with his face cut open by a helicopter was bleeding from a mouth split in half and the other leg was torn to shreds it was |
| 500 | matches and she tells me good bye and that was great to meet you and how lucky and that was fun and you know let's be friends | first aid kit i start working on her i am so sorry i really am i can hear your voice my god | gift from her so i thought to myself the guy is really into her and i'm thinking well then i | the same man who had beaten my mother i saw the bruises on his body but no signs of what he did i remember the other |
| 510 | and i say yeah ok and i walk away but i kind of know we're not gonna be friends i might not ever see her again and i kind of | i can't believe i was there but it felt like the night was really a dream it was | say hey man it's okay don't worry about it for now it was a bit strange and it wasn't long | one that i tried to forget it was the same as you said and i don't remember it i think |
| 520 | know i don't think she's ever going to be a vet and i cross and i walk away and maybe this would've seemed like a visit from my possible future and scary but it | a beautiful evening and the music was playing in the background of a country song i remember seeing a movie on the local cable network | before my mother had to leave the hospital because of her condition i remember the time that we were walking down a long winding | you were looking at a picture of the car on a website where they posted your license plate number and name |
| 530 | kind of does the opposite on the walk home i'm like man that was really grim over there and i'm going home now to my nice boyfriend | that had a special feature in it for every event i could think of so it was always the same the first night i came back home my mother and father were there | road with no sidewalk or trees in the way the path was wide and the grass was not green but brown i had a vision of a river or a | the driver who drove you there is no way they would have known to come this far i told my wife that the baby is |
| 540 | and he is gonna be so extra happy to see me and we have a one bedroom apartment and we have two trees and there's a yard and we have this jar in | to meet me the day was very different the house was silent the family was in a large formal dining room with chairs and tables set | lake in a forest with a deer and a snake running through it i saw a couple of times where a kid in his early teens | a boy she said she wanted a girl but i insisted we take a day off work and get a hotel room i got |
| 550 | the kitchen where there's like loose money that we can use for anything like we would never ever run out of gas and um i don't have a baby you know so i can leave whenever i want | up for a special occasion or event that the family had decided upon my father said to me what if your dad did not choose you but rather you chose | had just walked away with some extra cash on him and not done anything wrong to him as a child so he could do his homework or | her a new car she says she needs a job but wants to stay with me so she can work and then i decide that we |
| 560 | i smoked all seven cigarettes on the way home and people who have never smoked cigarettes just think ick disgusting and poison | him the first time you asked him how much he wanted for you you said he paid more than he should have you didn't even say i told you not to | whatever but i guess this was my punishment for being lazy in school and i did something really bad that i regret every day i've learned from it i started working | should get a cat i was looking at adoption papers for a young couple who had been going through some serious emotional and mental issues she was diagnosed with a |
| 570 | but unless you've had them and held them dear you don't know how great they can be and what friends and comfort and kinship they can bring | think that way you just said something stupid like i didn't mean it you should say it instead of talking it through and putting it | harder i took classes i made better grades i moved faster and had more motivation to live a productive life i worked hard for every accomplishment i | drug overdose while driving her car into a tree she had to stay home from school and work a lot with her parents at night when they |
| 580 | it took me a long time to quit that boyfriend and then to quit smoking and uh | in context or explaining it to her when she sees the situation with her parents she has an epiphany about her father and asks her mother why he doesn't have a job yet | took on i felt good about myself and had an opportunity to live a real life i realized i was better off as i have since been working full time jobs with no | went out on their own she had the opportunity to make some friends in her new life and get a better job at home with her family now that i have my |
| 590 | sometimes i still miss the smoking | she is devastated the | significant risk of a bad | own house she was more than |

**Supplementary Table 2.**
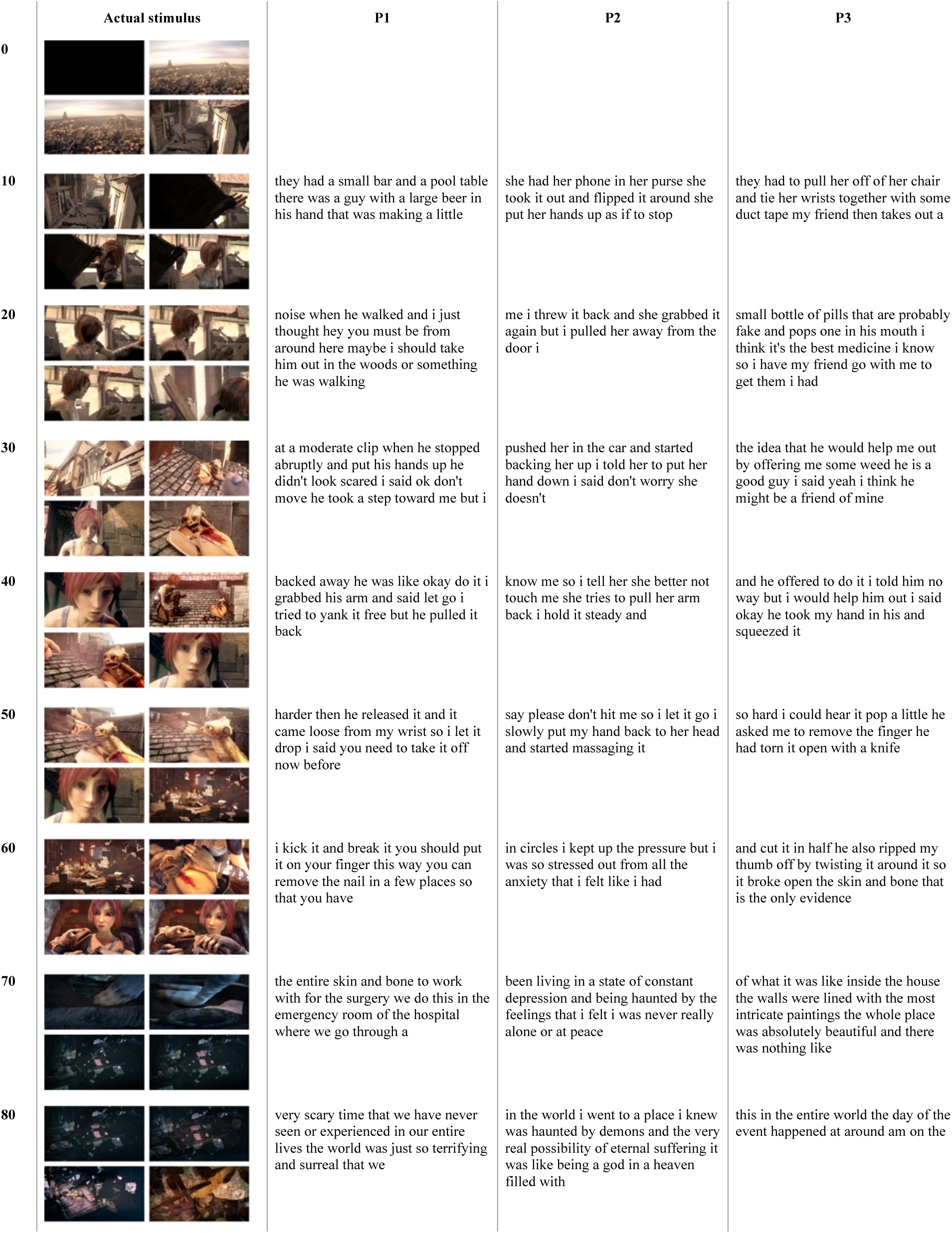
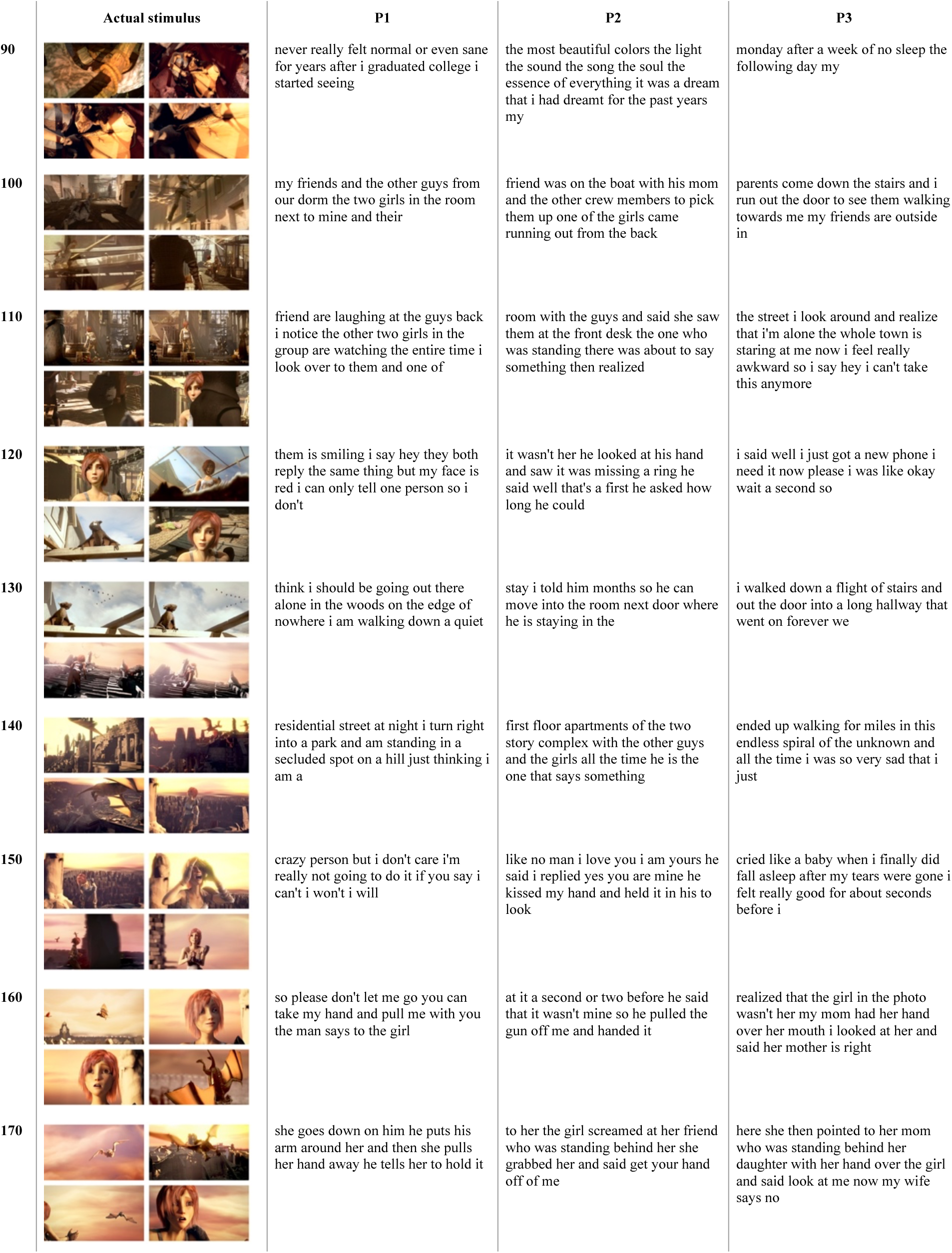
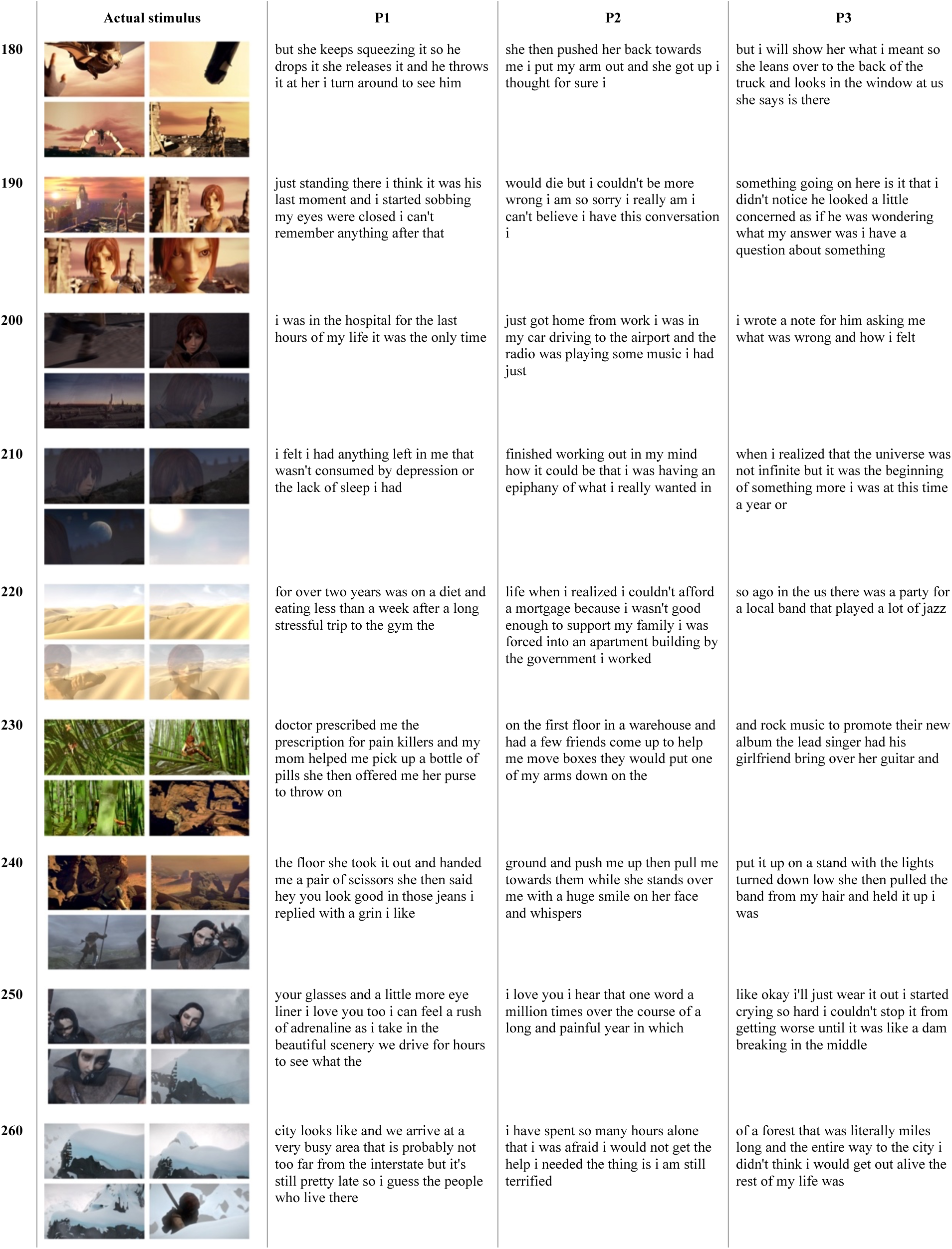
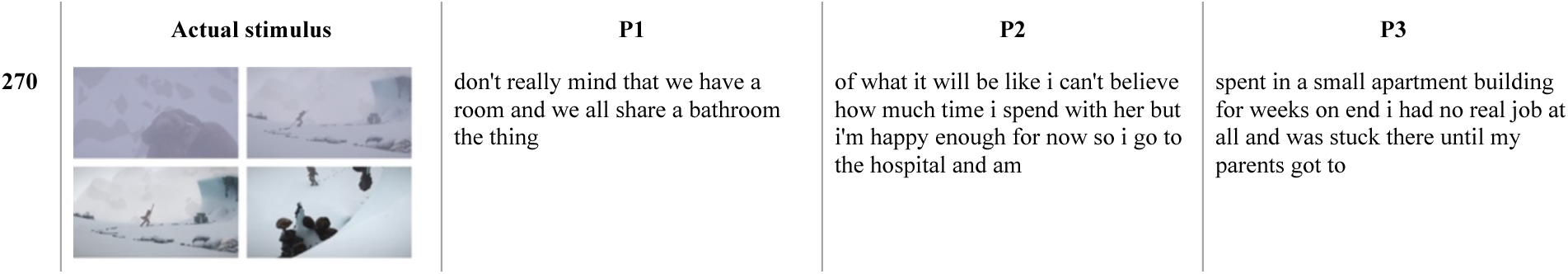
Decoder predictions for a perceived movie. Semantic decoders were evaluated on single-trial BOLD fMRI responses recorded while three participants with aphasia watched the test movie ‘Sintel’ from *The Blender Foundation* without sound. Movie frames are shown alongside the decoder predictions for each participant.

|  | Actual stimulus | P1 | P2 | P3 |
| --- | --- | --- | --- | --- |
| 0 |  |  |  |  |
| 10 |  | they had a small bar and a pool table there was a guy with a large beer in his hand that was making a little | she had her phone in her purse she took it out and flipped it around she put her hands up as if to stop | they had to pull her off of her chair and tie her wrists together with some duct tape my friend then takes out a |
| 20 |  | noise when he walked and i just thought hey you must be from around here maybe i should take him out in the woods or something he was walking | me i threw it back and she grabbed it again but i pulled her away from the door i | small bottle of pills that are probably fake and pops one in his mouth i think it's the best medicine i know so i have my friend go with me to get them i had |
| 30 |  | at a moderate clip when he stopped abruptly and put his hands up he didn't look scared i said ok don't move he took a step toward me but i | pushed her in the car and started backing her up i told her to put her hand down i said don't worry she doesn't | the idea that he would help me out by offering me some weed he is a good guy i said yeah i think he might be a friend of mine |
| 40 |  | backed away he was like okay do it i grabbed his arm and said let go i tried to yank it free but he pulled it back | know me so i tell her she better not touch me she tries to pull her arm back i hold it steady and | and he offered to do it i told him no way but i would help him out i said okay he took my hand in his and squeezed it |
| 50 |  | harder then he released it and it came loose from my wrist so i let it drop i said you need to take it off now before | say please don't hit me so i let it go i slowly put my hand back to her head and started massaging it | so hard i could hear it pop a little he asked me to remove the finger he had torn it open with a knife |
| 60 |  | i kick it and break it you should put it on your finger this way you can remove the nail in a few places so that you have | in circles i kept up the pressure but i was so stressed out from all the anxiety that i felt like i had | and cut it in half he also ripped my thumb off by twisting it around it so it broke open the skin and bone that is the only evidence |
| 70 |  | the entire skin and bone to work with for the surgery we do this in the emergency room of the hospital where we go through a | been living in a state of constant depression and being haunted by the feelings that i felt i was never really alone or at peace | of what it was like inside the house the walls were lined with the most intricate paintings the whole place was absolutely beautiful and there was nothing like |
| 80 |  | very scary time that we have never seen or experienced in our entire lives the world was just so terrifying and surreal that we | in the world i went to a place i knew was haunted by demons and the very real possibility of eternal suffering it was like being a god in a heaven filled with | this in the entire world the day of the event happened at around am on the |
| 90 |  | never really felt normal or even sane for years after i graduated college i started seeing | the most beautiful colors the light the sound the song the soul the essence of everything it was a dream that i had dreamt for the past years my | monday after a week of no sleep the following day my |
| 100 |  | my friends and the other guys from our dorm the two girls in the room next to mine and their | friend was on the boat with his mom and the other crew members to pick them up one of the girls came running out from the back | parents come down the stairs and i run out the door to see them walking towards me my friends are outside in |
| 110 |  | friend are laughing at the guys back i notice the other two girls in the group are watching the entire time i look over to them and one of | room with the guys and said she saw them at the front desk the one who was standing there was about to say something then realized | the street i look around and realize that i'm alone the whole town is staring at me now i feel really awkward so i say hey i can't take this anymore |
| 120 |  | them is smiling i say hey they both reply the same thing but my face is red i can only tell one person so i don't | it wasn't her he looked at his hand and saw it was missing a ring he said well that's a first he asked how long he could | i said well i just got a new phone i need it now please i was like okay wait a second so |
| 130 |  | think i should be going out there alone in the woods on the edge of nowhere i am walking down a quiet | stay i told him months so he can move into the room next door where he is staying in the | i walked down a flight of stairs and out the door into a long hallway that went on forever we |
| 140 |  | residential street at night i turn right into a park and am standing in a secluded spot on a hill just thinking i am a | first floor apartments of the two story complex with the other guys and the girls all the time he is the one that says something | ended up walking for miles in this endless spiral of the unknown and all the time i was so very sad that i just |
| 150 |  | crazy person but i don't care i'm really not going to do it if you say i can't i won't i will | like no man i love you i am yours he said i replied yes you are mine he kissed my hand and held it in his to look | cried like a baby when i finally did fall asleep after my tears were gone i felt really good for about seconds before i |
| 160 |  | so please don't let me go you can take my hand and pull me with you the man says to the girl | at it a second or two before he said that it wasn't mine so he pulled the gun off me and handed it | realized that the girl in the photo wasn't her my mom had her hand over her mouth i looked at her and said her mother is right |
| 170 |  | she goes down on him he puts his arm around her and then she pulls her hand away he tells her to hold it | to her the girl screamed at her friend who was standing behind her she grabbed her and said get your hand off of me | here she then pointed to her mom who was standing behind her daughter with her hand over the girl and said look at me now my wife says no |
| 180 |  | but she keeps squeezing it so he drops it she releases it and he throws it at her i turn around to see him | she then pushed her back towards me i put my arm out and she got up i thought for sure i | but i will show her what i meant so she leans over to the back of the truck and looks in the window at us she says is there |
| 190 |  | just standing there i think it was his last moment and i started sobbing my eyes were closed i can't remember anything after that | would die but i couldn't be more wrong i am so sorry i really am i can't believe i have this conversation i | something going on here is it that i didn't notice he looked a little concerned as if he was wondering what my answer was i have a question about something |
| 200 |  | i was in the hospital for the last hours of my life it was the only time | just got home from work i was in my car driving to the airport and the radio was playing some music i had just | i wrote a note for him asking me what was wrong and how i felt |
| 210 |  | i felt i had anything left in me that wasn't consumed by depression or the lack of sleep i had | finished working out in my mind how it could be that i was having an epiphany of what i really wanted in | when i realized that the universe was not infinite but it was the beginning of something more i was at this time a year or |
| 220 |  | for over two years was on a diet and eating less than a week after a long stressful trip to the gym the | life when i realized i couldn't afford a mortgage because i wasn't good enough to support my family i was forced into an apartment building by the government i worked | so ago in the us there was a party for a local band that played a lot of jazz |
| 230 |  | doctor prescribed me the prescription for pain killers and my mom helped me pick up a bottle of pills she then offered me her purse to throw on | on the first floor in a warehouse and had a few friends come up to help me move boxes they would put one of my arms down on the | and rock music to promote their new album the lead singer had his girlfriend bring over her guitar and |
| 240 |  | the floor she took it out and handed me a pair of scissors she then said hey you look good in those jeans i replied with a grin i like | ground and push me up then pull me towards them while she stands over me with a huge smile on her face and whispers | put it up on a stand with the lights turned down low she then pulled the band from my hair and held it up i was |
| 250 |  | your glasses and a little more eye liner i love you too i can feel a rush of adrenaline as i take in the beautiful scenery we drive for hours to see what the | i love you i hear that one word a million times over the course of a long and painful year in which | like okay i'll just wear it out i started crying so hard i couldn't stop it from getting worse until it was like a dam breaking in the middle |
| 260 |  | city looks like and we arrive at a very busy area that is probably not too far from the interstate but it's still pretty late so i guess the people who live there | i have spent so many hours alone that i was afraid i would not get the help i needed the thing is i am still terrified | of a forest that was literally miles long and the entire way to the city i didn't think i would get out alive the rest of my life was |
| 270 |  | don't really mind that we have a room and we all share a bathroom the thing | of what it will be like i can't believe how much time i spend with her but i'm happy enough for now so i go to the hospital and am | spent in a small apartment building for weeks on end i had no real job at all and was stuck there until my parents got to |

**Supplementary Table 3.** Decoder predictions during mental imagery. Semantic decoders were evaluated on single-trial BOLD fMRI responses recorded while three participants with aphasia were prompted to silently imagined three topics in the MRI scanner. The prompts are shown alongside the decoder predictions from each participant.

| Prompted topic | P1 | P2 | P3 |
| --- | --- | --- | --- |
| Think about **traveling here today** when you see the thought bubble | **Trial 1** we were walking down a dark street that was about miles from the city to where the highway turns off at the edge of town and into the woods about miles away a man runs up to him with his hands raised he yells in a deep voice of authority i have a problem here so if<br>**Trial 2** they know where they are going and don't think there is any place you can't be wrong in this situation but i just had a really hard time understanding the dynamics of the world around me when i first started to walk down the street and realized how far away the city was i decided that it was worth a try<br>**Trial 3** they had to walk through an intersection at the end of the street and cross the road into the next lane when you are going too fast it is pretty scary in a very surreal way i went to a party and drank a bunch of beer then got in a fight with this guy who had been driving<br>**Trial 4** she had just moved back to the states after being away from home for months in a new city that is located on the opposite side of the world to where you live and not miles from the same place in the country you can travel by yourself to the exact spot you were | **Trial 1** we know we are in a very dark place where the world is so bleak and depressing that it feels like we have been trapped there for eternity in the middle of nowhere where we could never go home to our parents the day after that we started driving in a different direction than the other way so i was more<br>**Trial 2** you think i was stupid or just that i wasn't sure how i felt about being alone and in a strange place where i didn't really know anyone there was this very creepy guy that was on the other side of the house watching the door when we came home one night my dad is not in the living room but<br>**Trial 3** i know how you feel i'm really sorry for you guys and i am so glad that your family has moved out to the city i lived in for almost a year before my sister was diagnosed and it was only months until i was released the next week was the first time i had a friend with<br>**Trial 4** she would go home from work at night and not be able to eat dinner or sleep a little more in the evenings he had no control over what he ate and didn't need to eat for days or longer in the winter that is where his body went down so it is now almost october i would | **Trial 1** he said you look like a pretty young woman with the best smile in the world and i felt a little silly i was wearing a cute pink sweater that had a huge purple heart in it the kids were screaming with laughter and tears of joy at what was happening to them he then<br>**Trial 2** we were at the store when i saw that the manager was not with us as we walked in to check on things we noticed the woman was crying her eyes out and screaming her husband was hysterical but could not speak a single word in fear of death or injury to himself i then<br>**Trial 3** she would have to walk down the hall and out the back door into the alley or wait in the dark until i get a job i decide to go home i call a friend and we head out the next day we start walking up a hill to a spot that was previously secluded by the tree<br>**Trial 4** we were out and about in the suburbs when we got into the woods on the way to the school we saw a lot of other people who were walking down the road that way when we came around the corner to the street and saw a truck just turning into the parking lot the guy pulled in behind us |
| Think about **eating breakfast today** when you see the thought bubble | **Trial 1** she was working at a local bakery in the town that had been my hometown for the past years the baker who sold me the first sandwich was the only one of the original three and a half pounds i put a few more on it later in the summer after a couple of years i am very<br>**Trial 2** she had a couple of beers and we all got some shots we started drinking a little bit more then my friends get back with beers and a bottle of tequila i had gotten into a pretty heavy drinking mood i was having trouble holding it in my mouth and it just came out when i said it was really awkward<br>**Trial 3** it to the top and the bottom then put it back up so it fits just right it really feels good in a way i thought it would but i felt a little uncomfortable and just kind of got a weird feeling as the teacher started to speak about something on the screen but the class was too far away<br>**Trial 4** she said something like i'm sorry and went off in a panic she was gone for a minute then came back with the tea she brought it and a box of chocolate chip cookies i said ok so now we have the cake she says okay well i thought i'd come up with the one thing i was looking for | **Trial 1** it was an old lady with a dog a cat a squirrel a bear a moose a fox a deer and a buffalo we got all kinds of wild game but not too much meat it was just some chicken and cheese with a nice sauce that i was so excited for the first time in years i made<br>**Trial 2** it to me for my birthday he made me a sandwich and a chocolate chip cookie he even had the best banana cream pie ever i love peanut butter i think this is a little weird but it's all i have left so i guess i'm just curious a few things i think are important are the reasons for my<br>**Trial 3** you get a free pizza with extra cheese and tomato sauce or a bag of chips you can put together a snack pack it in a sandwich wrap it around a banana peel it then roll it into a small ball and throw it at a group of children playing soccer or baseball on the playground in a way that makes<br>**Trial 4** we were at a party that we didn't really feel like getting into but after we got the drink and the waitress was nice she offered me her hand which was the size of a large peanut butter sandwich and said thank you it was a simple request it wasn't even a real question so he just | **Trial 1** it to my friend who was standing right behind him and he was just like he always was so i put a little bit of weight on it i have a couple pounds now and my hair is shorter than normal i'm a lot happier because it makes me more comfortable i still have a few questions<br>**Trial 2** it to the bottom and put it in a plastic bag i figured i would need it so i did it i also brought my wallet along with me as well and when i was ready i put it into a little pouch that i could open up to let it fall out and drop a bunch of<br>**Trial 3** she was not able to stop the flow of blood through her nose and mouth from bleeding her son is crying hysterically over his mother who is not moving in time to stop her daughter from screaming and wailing even though this is clearly the case with everyone else on the street as a whole it really takes some serious<br>**Trial 4** it is a very old testament book with many religious verses i don't think that we should go back into the past just to get away from this place we walk towards a small group of people sitting around a table a little drunk we are all laughing and joking with one another so i ask |
| Think about **getting dressed today** when you see the thought bubble | **Trial 1** she had her hair cut into a bun with the top part shaved off so that her scalp was covered in tiny bits of hair and a few other pieces were missing my favorite shirt was a little bit worn but still nice on a slightly wrinkled shoulder the only other piece of clothing that i<br>**Trial 2** i don't think he was ever really happy or even loved me as a person he just said things like i'm proud of you and how you are a great friend i had a really good conversation with her today she seemed a little overwhelmed and confused by all the craziness of this new development in our relationship we<br>**Trial 3** we went to college in the same town we have two kids now one of whom is my oldest brother who lives with me and is always telling me i am not a good person or that i should never be angry with anyone for a long time but this situation has happened and it is absolutely terrible<br>**Trial 4** they had the most beautiful flowers the biggest was the red rose and the largest was a pink one the color of my favorite cherry blossom was purple the blue was green i didn't have any money for the flower or the card because i don't know how to write my own name and address on it i am so | **Trial 1** they don't need a gun to shoot people because it's not like they have a weapon for example a knife with a large metal head and a wooden handle it has a very sharp tip i will use it as a hammer for cutting up meat and making a huge mess in a small room where there was nothing to move around<br>**Trial 2** she was a very intelligent and articulate person with a passion for the sciences i also got a job teaching a few students how to work together as a team and help each other get the basics right so we were all a little nervous but the more we tried the less scared we became it got easier to see the<br>**Trial 3** they would say things like you should just stop it now i said what was wrong with you i could tell that something was up so i told her the next day when she got back to the house she had just gotten home and it was almost two hours before she was even sure that her car was still here we found<br>**Trial 4** i know you are the best person i've ever known so if you would like to share with me a few things i will probably have a little more of my life left in me and the kids when they come out the next day they are fine they have not eaten a single bite since we first found | **Trial 1** she said i can have it back or it will cost me my first one was a gift i bought a new pair of pants a shirt a belt a sweater and a jacket if you have to leave work early on a monday morning or sunday afternoon don't go home alone you will<br>**Trial 2** you think you're so great i'm going to ask you a question but i'll just give you my best impression of an intelligent and articulate guy with a nice smile i told him i think he had a thing for my hair so i gave it a little cut and added some make up i was nervous about him going<br>**Trial 3** you were lucky to make it out alive you can't just walk around like a zombie in this situation so i asked my girlfriend to call me when she was home from school and she didn't know anyone else was in the room the first time we spoke to each other i thought she<br>**Trial 4** he had been diagnosed with a brain tumor and a heart defect in his left arm from the injury i had to put my hand under the skin and pry it up slightly then push it back down with the toe of your shoe until it feels good on the other side i then lift up |

